# Differential predictive value of resident memory CD8^+^T cell subpopulations in non-small-cell lung cancer patients treated by immunotherapy

**DOI:** 10.1101/2024.03.07.583820

**Authors:** Léa Paolini, Thi Tran, Stéphanie Corgnac, Jean-Philippe Villemin, Marie Wislez, Jennifer Arrondeau, Ludger Johannes, Jonathan Ulmer, Louis-Victorien Vieillard, Joséphine Pineau, Alain Gey, Valentin Quiniou, Pierre Barennes, Hang Phuong Pham, Nadège Gruel, Milena Hasan, Valentina Libri, Sébastien Mella, Sixtine De Percin, Pascaline Boudou-Rouquette, Isabelle Cremer, Hélène Blons, Karen Leroy, Pierre Laurent-Puig, Hortense De Saint Basile, Laure Gibault, Patrice Ravel, Fathia Mami- Chouaib, François Goldwasser, Elizabeth Fabre, Diane Damotte, Eric Tartour

## Abstract

A high density of resident memory T cells (T_RM_) in tumors correlates with improved clinical outcomes in immunotherapy-treated patients. However, in preclinical models, only some subpopulations of T_RM_ are associated with cancer vaccine efficacy.

We identified two main T_RM_ subpopulations in tumor-infiltrating lymphocytes derived from non-small cell lung cancer (NSCLC) patients: one co-expressing CD103 and CD49a (DP), and the other expressing only CD49a (MP); both exhibiting additional T_RM_ surface markers like CD69. DP T_RM_ exhibited greater functionality compared to MP T_RM_. Analysis of T-cell receptor (TCR) repertoire and of the stemness marker TCF-1 revealed shared TCRs between populations, with the MP subset appearing more progenitor-like phenotype. In two NSCLC patient cohorts, only DP T_RM_ predicted PD-1 blockade response. Multivariate analysis, including various biomarkers (CD8, TCF1^+^CD8^+^T cells, and PD-L1) associated with responses to anti-PD(L)1, showed that only intra-tumoral infiltration by DP T_RM_ remained significant. This study highlights the non-equivalence of T_RM_ populations and emphasizes the importance of distinguishing between them to better define their role in antitumor immunity and as a biomarker of response to immunotherapy.

## Introduction

More than two decades ago, pioneering work on vesicular stomatitis virus and *Listeria monocytogenes* revealed the presence and persistence of noncirculating resident memory T cells (T_RM_) in nonlymphoid organs after the resolution of the primary infection (Masopust, Vezys et al., 2001). It rapidly became apparent that these T_RM_ constituted a specific lineage associated with a profile of transcription factors including Blimp1, Runx3, and Notch family proteins (Hombrink, Helbig et al., 2016). In terms of their phenotype, these T_RM_ express core markers such as CD69, CD103, and CD49a, together with the loss of expression of other markers such as CD62L, CCR7, S1PR1, and KLF2, favoring the persistence of these cells within tissues (Kumar, Ma et al., 2017, Mami-Chouaib, Blanc et al., 2018). T_RM_ have innate-like “sensing and alarming” properties that enable them to recruit other immune cells to control microbial infections (Ariotti, Hogenbirk et al., 2014, Ge, Monk et al., 2019, Schenkel & Masopust, 2014). As they are located at the site of inflammation in the tissues, T_RM_ respond much more rapidly to reinfection and provide superior protection compared to circulating memory cells, including central memory and effector memory T cells (Ariotti et al., 2014, Iijima & Iwasaki, 2014, Schenkel & Masopust, 2014).

In a range of preclinical cancer models, we have shown that T_RM_ are required for the efficacy of anti-tumor vaccines against mucosal tumors such as lung and head and neck cancer (Nizard, Roussel et al., 2017, Sandoval, Terme et al., 2013). T_RM_ can also activate dendritic cells (DCs) to increase the numbers of tumor-specific CD8^+^ T cells, conferring protection against tumor rechallenge (Menares, Galvez-Cancino et al., 2019). In humans, high levels of intratumoral T_RM_ infiltration have been associated with better clinical outcomes in multiple solid tumors including lung, melanoma, bladder, breast, cervical, ovarian, endometrial, gastric, and colorectal cancers receiving standard-of-care treatments (Djenidi, Adam et al., 2015, Ganesan, Clarke et al., 2017, Nizard et al., 2017, Okla, Farber et al., 2021, Wang, Wu et al., 2015). More recently in humans, correlative studies in non-small cell lung cancer (NSCLC), bladder cancer, and melanoma have revealed an association between tumor infiltration by CD8^+^ T cells with a resident phenotype before immunotherapy and responses to immune checkpoint blockade (Attrill, Owen et al., 2022, Banchereau, Chitre et al., 2021, Corgnac, Malenica et al., 2020, Zitti, Hoffer et al., 2023).

To explain this predictive role of T_RM_ during immunotherapy, various groups have shown that during neoadjuvant treatment in breast and head and neck cancer, CD8^+^ tumor-infiltrating lymphocytes (TILs) with a tissue-resident phenotype expand and are characterized by a gene expression program related to activation, cytotoxicity, and effector functionality (Bassez, Vos et al., 2021, Luoma, Suo et al., 2022). The same expansion of CD8^+^ T_RM_ has been observed after anti-PD-1/PD-L1 monotherapy or combined anti-CTLA-4 treatment in melanoma, lung, breast, and esophageal cancer (Corgnac et al., 2020, Edwards, Wilmott et al., 2018, Jaiswal, Verma et al., 2022, Virassamy, Caramia et al., 2023). The role of T_RM_ as immunotherapeutic targets raises the possibility that other effectors recruited secondarily or present in the blood may play a role in the efficacy of immunotherapy (Yost, Satpathy et al., 2019).

Given the anti-tumor role of T_RM_ and their prognostic and predictive value in the context of patient responses to immunotherapy, the issue of optimal strategies for inducing or increasing this population is becoming a major challenge in immuno-oncology. In preclinical models, we have shown that the nasal route, but not the muscular route, induces T_RM_ with a CD103^+^CD49a^+^CD69^+^ phenotype (Nizard et al., 2017, Sandoval et al., 2013). This induction was associated with the inhibition of the growth of lung or head and neck tumors. In infectious and oncological models, other groups have also documented a correlation between the preferential ability of mucosal immunization to induce T_RM_ expansion and protection against the development of cancers and viral infections (Jeyanathan, Fritz et al., 2022, Stewart, Counoupas et al., 2022, Sun, Peng et al., 2016). However, other studies based on vaccinations using systemically administered recombinant viral vectors have shown that T_RM_ can be induced in the lungs (Douguet, Fert et al., 2023). This result may be explained by the use of viruses that enable vaccine dissemination in the pulmonary or head and neck compartments. However, mRNA-based vaccinations have also been shown to induce T_RM_ when administered via the intramuscular (i.m.) route, but the cells induced in this context generally express CD69 or CD49a without any concomitant CD103 expression (Kunzli, O’Flanagan et al., 2022). The fact that different sub-populations of T_RM_ exist with differing core marker (CD103, CD49a, CD69) expression patterns has been reported in various tissues, and this effect has sometimes been linked to the different functions of these cells (Cheuk, Schlums et al., 2017). Although CD103 is considered a hallmark of T_RM_, persistent CD103-negative T_RM_ have also been described in tissues. In contrast with CD103+ T_RM_, these cells were able to develop in a TGFβ-independent manner (Bergsbaken & Bevan, 2015, Schenkel, Fraser et al., 2014, Steinert, Schenkel et al., 2015).

In this work, we aimed to better characterize these different T_RM_ subpopulations in mice and humans and to determine whether they play distinct roles as predictors of responses to immunotherapy in patients with NSCLC.

## RESULTS

### 1. Different immunization routes give rise to distinct subpopulations of resident memory CD8^+^T lymphocytes

Our previous work had shown that only the intranasal (i.n) immunization route induced resident memory CD103-expressing CD8^+^T cells in bronchoalveolar lavage fluid (BAL), and this mucosal immunization route was associated with tumor rejection (Nizard et al., 2017). In recent years, it has emerged that there are different T_RM_ subpopulations defined by the markers CD103, CD49a, and CD69 (Mami-Chouaib et al., 2018). Only i.n. vaccination with the STxB-E7 vaccine combined with αGalCer induced D^b^-restricted E7_49-57_ peptide-specific CD8^+^T cells co-expressing CD103 and CD49a in the BAL (Fig. 1A). In contrast, the i.m. route also induced E7-specific CD8^+^ T cells expressing CD49a but not CD103 (Fig. 1A). Both CD103^+^CD49a^+^ and CD103^neg^CD49a^+^CD8^+^ T cell populations induced by the i.n route expressed high levels of CD69 (For CD103^+^CD49a^+^ : Mean 94.82% ± 1.86% and for CD103^neg^CD49a^+^ Mean: 95.5% ± 1.49%) (Fig. 1A, lower right), whereas the frequency of CD8^+^ T cells specific for E7 and expressing neither CD103 nor CD49a, which are considered to be effector T cells, exhibited weaker CD69 expression (Mean 49.24% ± 10.24%)(Fig. 1A, lower right). Immunization via the i.n. route induced a marginal CD103^+^CD49a^−^T cell population (< 5%)(Fig. 1)

**Fig 1:**
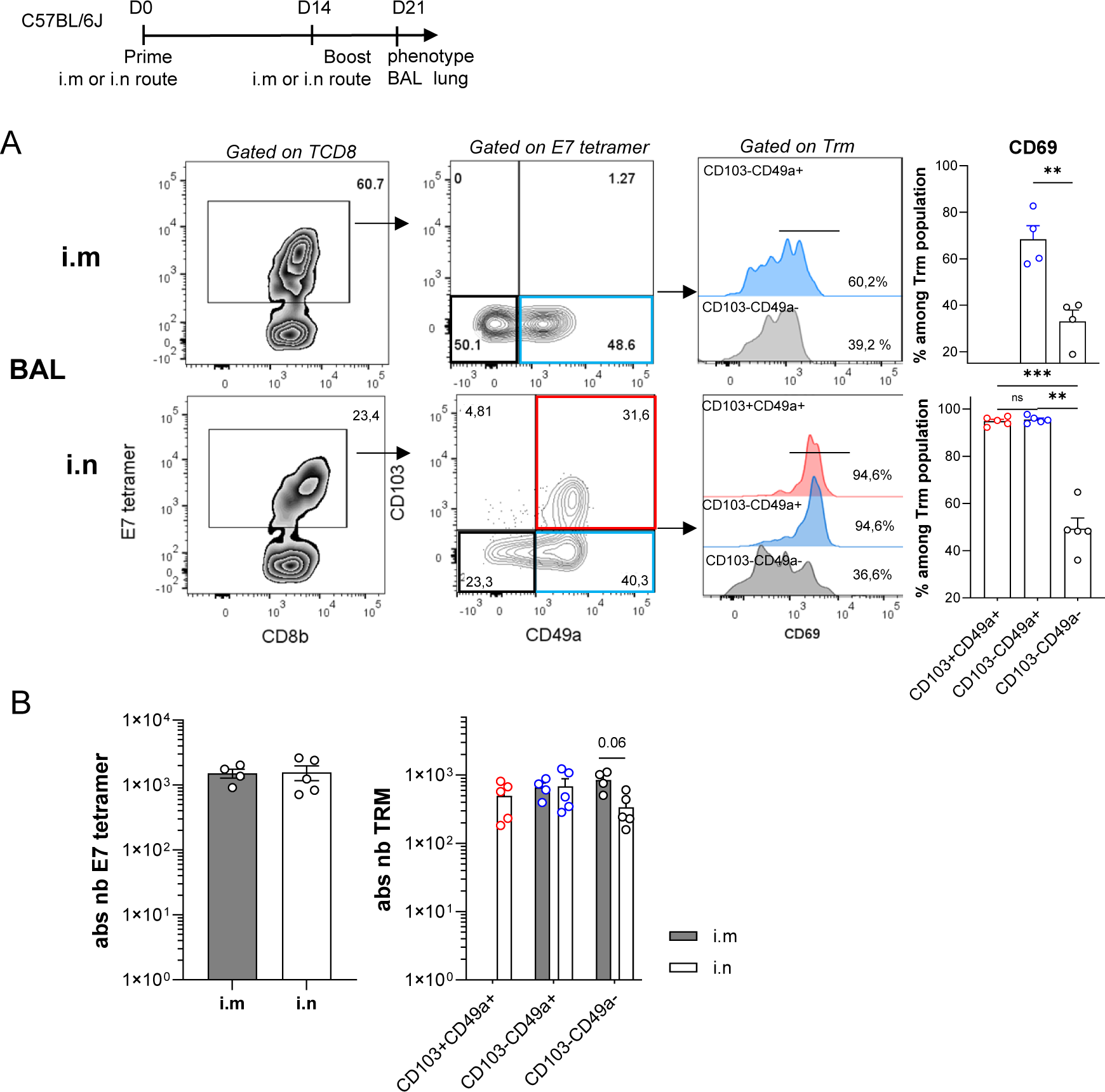
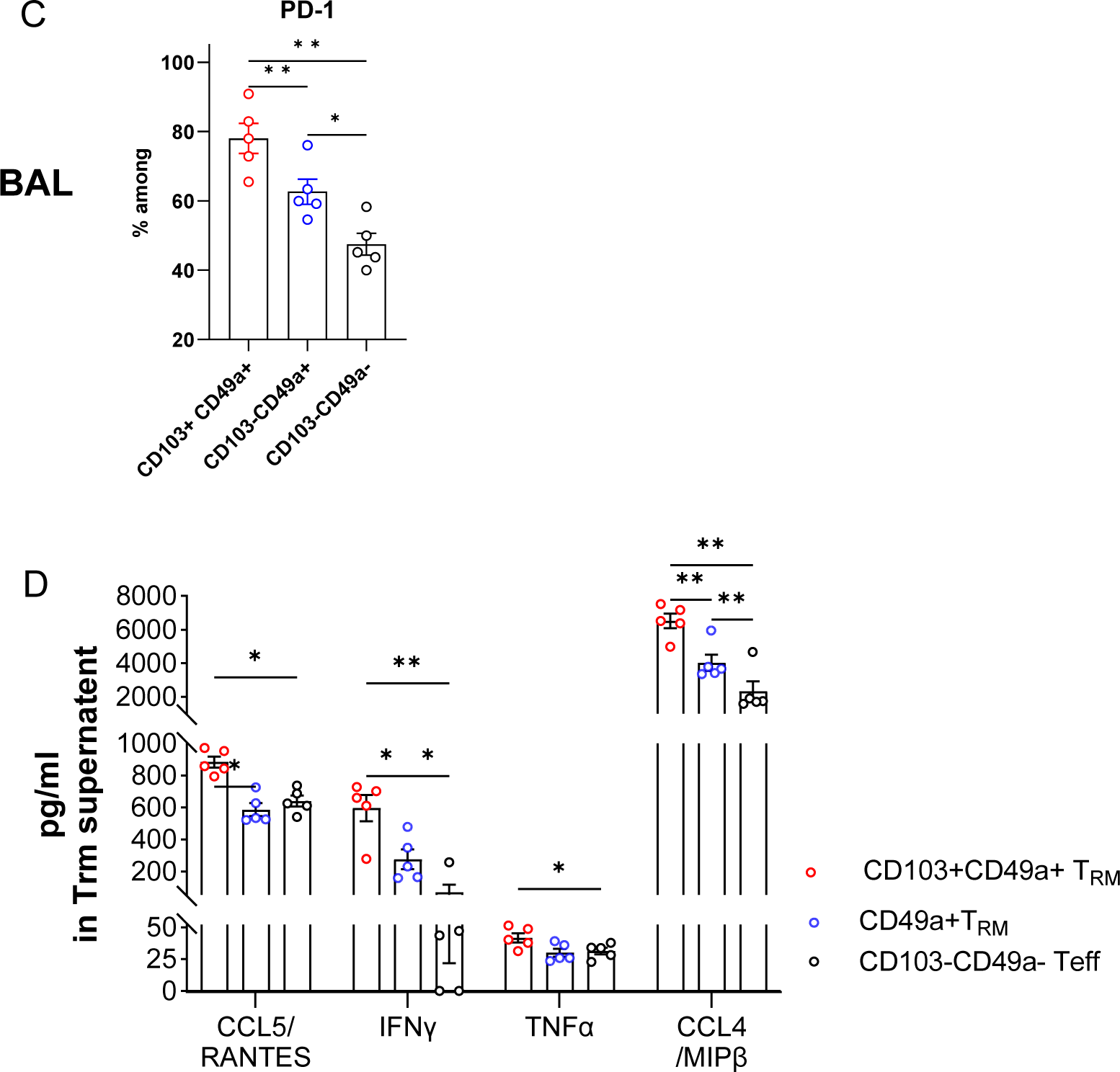
Different immunization routes give rise to distinct subpopulations of resident memory CD8^+^T lymphocytes with different phenotype and functionality. C57BL/6J mice were immunized with STxB-E7 and aGalcer intranasally (i.n.) or intramuscularly (i.m.) on days 0 and 14, and were sacrificed on day 21. CD8a APCefluo780 (5 µg) was injected (i.v.) 5 min before sacrifice to discriminate between circulating CD8 cells and resident CD8 cells. (A) Representative flow cytometry plots in BAL (bronchoalveolar lavage fluid) of E7 tetramer-specific CD103^+^CD49a^+^ T_RM_, CD103^−^CD49a^+^ T_RM_, and CD103^−^CD49a^−^ Teff, and the frequency of CD69 expression among these populations. (B) Absolute numbers of (left) E7-tetramer-specific CD8^+^ and (right) CD103^+^CD49a^+^ T_RM_, CD103^−^CD49a^+^ T_RM_, and Teff CD103^−^CD49a^−^ in the BAL. Data are means ± SD. One representative experiment with 3-5 mice from 2 independent experiments is shown. Data were compared between groups via two-sided Mann-Whitney t-tests. (C) Percentage of PD1 expression among E7-specific CD103^+^CD49a^+^ T_RM_, CD103^−^CD49a^+^ T_RM_, and CD103^−^CD49a^−^ Teff in the BAL. (D) E7-specific CD103^+^CD49a^+^ T_RM_, CD103^−^CD49a^+^, and CD103^−^CD49a^−^ Teff were sorted from the BAL on day 21 and stimulated (10,000 cells/well) with E7_49-57_ peptide (10 µg/ml) for 18 h. Then, supernatants were harvested and multiplex cytokine detection was performed. Data are means ± SD. One representative experiment with 3-5 mice from 2 independent experiments is shown. Data were compared within groups via paired one-way ANOVAs with Tukey’s multiple comparisons test. *P<0.05, **P<0.01, ***P<0.001, ****P<0.0001

It should be noted that i.m.-induced CD49a^+^CD103^−^CD8^+^T cells were less likely to express CD69 as compared to i.n.-induced ones (Mean: 68.4% ± 11.6%)(Fig. 1A, upper right), but their numbers were equivalent in the BAL irrespective of the immunization route (Fig. 1B). These results were replicated by analyzing the same resident memory CD8^+^ T cell subpopulations in lung parenchyma after i.m or i.n vaccination (Fig. S2 A,B).

Similarly, using another vaccine system consisting of the protein ovalbumin mixed with the adjuvant c-di-GMP, only immunization via the i.n. route induced ovalbumin-specific CD103-expressing CD8^+^T cells in the BAL (Sup Fig. 2C,D). The subcutaneous route, like the i.m. route above, induced only CD103^neg^CD49a^+^CD8^+^T cells (Fig. S2 C,D). These two T_RM_ populations expressed more granzyme B than effector T cells (Fig. S4 and data not shown). This difference was significant for the CD103^+^CD49a^+^ population.

**Figure 2:**
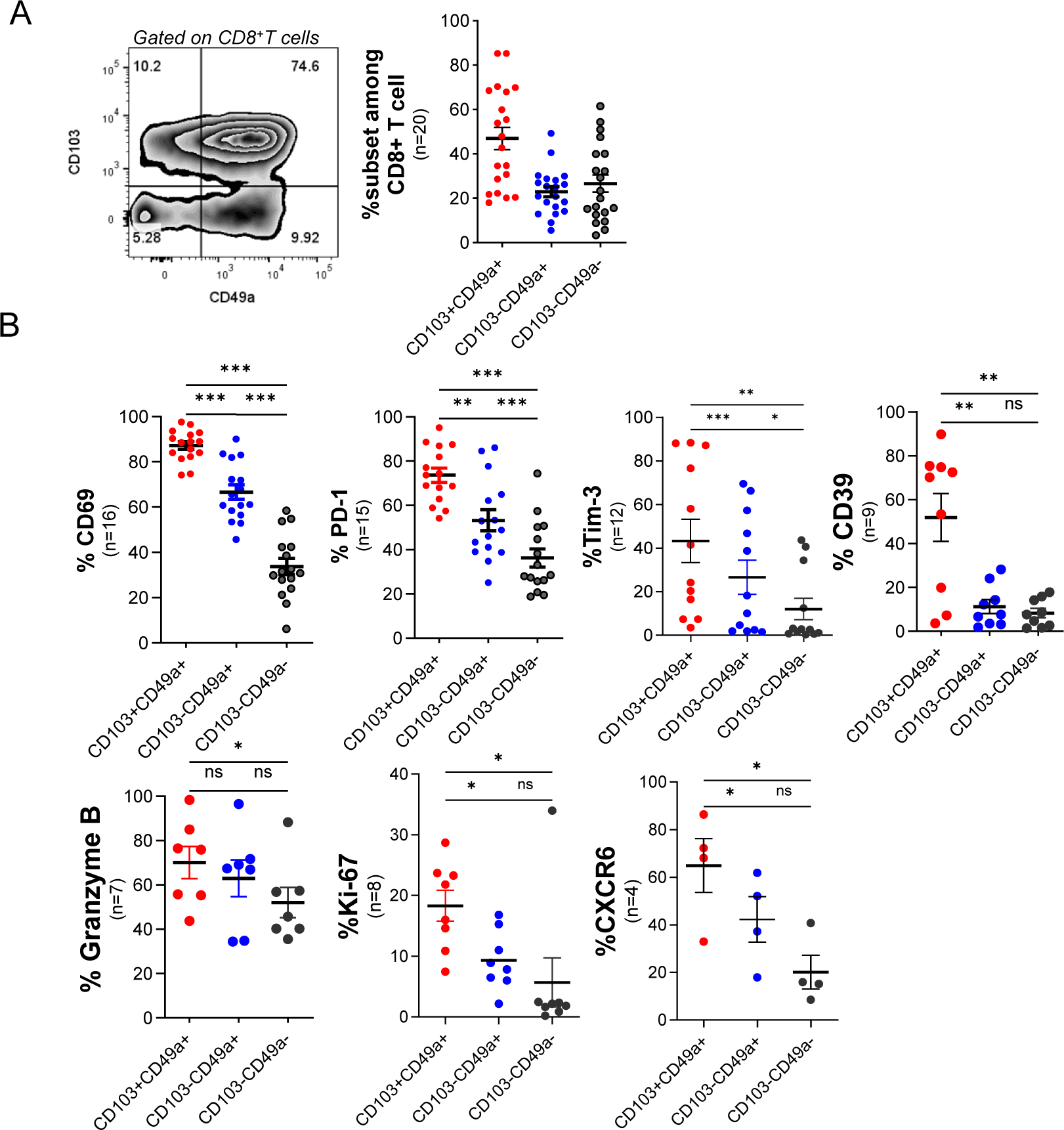
Phenotypic analyses of subpopulations of T_RM_ and effector T cells among TILs derived from lung cancer patients. Fresh biopsies from lung cancer patients (n = 20) were dissociated and digested, and flow cytometry was used to analyze TILs. The number of TILs tested per marker is shown below each figure. (A) The percentages of T_RM_ subpopulations (CD103^+^CD49a^+^ and CD49a^+^CD103^+^) among CD8^+^T cells, as well as non-effectors T_RM_ (CD49a^−^CD103^−^) among CD8^+^T cells are shown. (B) The percentages of different markers defining T_RM_ (CD69, CXCR6), exhausted T cells (PD^−^ 1, Tim-3, CD39), cytotoxicity (GZMB), and proliferation (Ki67) are shown among the two populations of T_RM_ and non-T_RM_ effectors (CD103^−^CD49a^−^). Significance was determined using paired t-tests. P < 0.05 was regarded as statistically significant. *P < 0.05, **P < 0.01, ***P < 0.001; n = 4-20

Characterization of these resident memory E7-specific CD8^+^ T cell subpopulations (CD103^+^CD49a^+^ and CD103^neg^CD49a^+^) revealed that the CD103^+^CD49a^+^CD8^+^T cells expressed more PD-1 (Mean: 78.04% ± 9.65%) as compared to the CD103^neg^CD49a^+^ population (Mean 62.66% ± 8.14%) and the CD103^neg^CD49a^neg^ effector CD8^+^ T cell population (Mean: 47.46% ± 7.03%). (Fig. 1C). Interestingly, the resident memory CD8^+^ T cell population co-expressing CD103 and CD49a appeared to be more functional after antigenic stimulation with a peptide derived from the E7 protein, secreting more IFNγ, TNFα, CCL4, and CCL5 (Fig. 1D) highlighting the more functional and protective phenotype of these cells.

### 2. Distinct subpopulations of T_RM_-type CD8^+^T lymphocytes co-exist among lung TILs

TILs were obtained from 20 dissociated tumors from patients with NSCLC. The same T_RM_ subpopulations identified in mice (CD103^+^CD49a^+^ and CD103^−^CD49a^+^) were detected in humans, with the CD103^+^CD49a^+^ population being present at a higher frequency (Mean: 47% ± 22.52% of total CD8)(Fig. 2A). Nearly all of these T_RM_ exhibited an effector memory (EM) T cell phenotype (CCR7^−^CD45RA^−^) (Fig. S3). The two T_RM_ subpopulations expressed CD69 at a higher frequency (Mean %CD69^+^: 87.2% ± 6.9% in CD103^+^CD49a^+^ and 66.6% ± 12.8% in CD103^neg^CD49a^+^) as compared to the effector CD8^+^T cell population (CD103^neg^CD49a^neg^; 33.8% ± 14.12%) (Fig. 2B). The CD103^+^CD49a^+^ cells expressed exhaustion markers (PD-1, Tim-3, CD39) and the proliferation marker Ki67 significantly more frequently (Fig. 2B). These data were confirmed through single-cell transcriptomic analyses of intratumoral CD8^+^ T cells, revealing that the resident memory CD103^+^CD49a^+^CD8^+^ T cells exhibited higher levels of exhaustion and proliferation marker expression relative to the CD103^−^CD49a^+^CD8^+^ T cell population (Fig. S4).

We regard the CD103^neg^CD49a^+^ population as a T_RM_ population, as most of these cells express CD69, which is considered to be a T_RM_ marker (Kumar et al., 2017). They also express the transcription factors Hobit and Runx3, which are hallmarks of T_RM_. However, in contrast to mice, these transcription factors are not enriched in human T_RM_ (Fig. S5) (Hombrink et al., 2016).

### 3. Relationship between the T_RM_ subpopulations in terms of differentiation

Four fresh tumors were used to conduct single-cell transcriptomic analyses, revealing that, among all the patients, the five most frequent clonotypes (TRA or TRB) of the double positive T_RM_ CD8^+^T cell subpopulation (CD103^+^CD49a^+^) were also present in the single positive T_RM_ CD8^+^ T cell subpopulation (CD103^neg^CD49a^+^) as well as in the CD103^neg^CD49a^neg^ effector CD8^+^ T cell population (Fig. 3A). These clonotypes were most amplified in the CD103^+^CD49a^+^CD8^+^ T cells, followed by the CD103^neg^C49a^+^CD8^+^T cell population and finally the CD103^neg^CD49a^neg^ CD8^+^T cell population (Fig. 3B), suggesting possible differentiation and proliferation of effector CD8^+^ T lymphocytes into the resident memory CD103^neg^CD49a^+^CD8^+^ T cells and then into resident memory CD103^+^CD49a^+^CD8^+^ T cells. Based on the Jaccard and the Morisita-horn dissimilarity indices (Fig. 3C and 3D, respectively), we found that for 3 out of 4 patients analyzed, closer repertoire composition and less dissimilarity (index close to 0) was observed between the CD103^+^CD49a^+^ and the CD103^neg^CD49a^+^ CD8^+^T cell populations as compared to the CD103^neg^CD49a^neg^ population, reinforcing the relationship between the two T_RM_ populations. With respect to the trajectories of these three subpopulations, another argument supports a more terminal differentiation of the CD103^+^CD49a^+^ T_RM_ population since they express less TCF1 progenitor (Fig. Sup 6). Indeed, in mice, the % of cells expressing TCF-1, a stemness marker, is lower (Mean 57.03% ± s.d. 6.37%) in the CD103^+^CD49a^+^ CD8^+^T cell population, than in the CD103^neg^CD49a^+^CD8^+^T cells (Mean 73.1% ± 5.45%) and in the CD103^neg^CD49a^neg^ effector CD8^+^T populations (Mean 71.78% ± 9.45%)(Fig. S6). In humans, multiplex *in situ* immunofluorescence imaging revealed that the CD103^+^CD49a^+^CD8^+^T cell population does not express TCF-1 (results not shown), as described previously (Wu, Madi et al., 2020).

**Figure 3.**
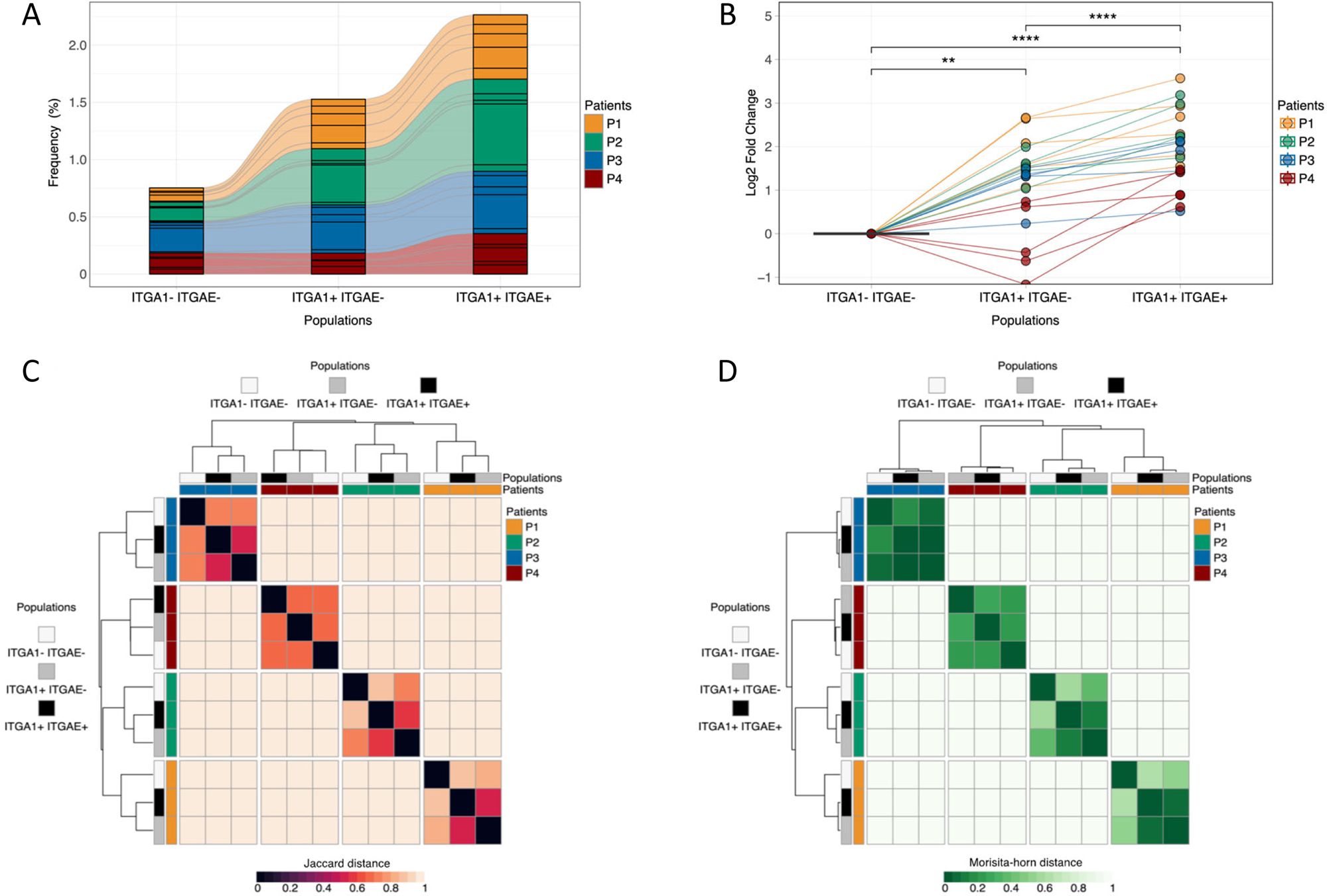
TCR sharing among the subpopulations of resident memory CD8^+^T cells. (A) Tracking of the most predominant clonotypes within the ITGA1^+^(CD49a)/ITGAE^+^ (CD103) population. Alluvial plots represent the relationships between the frequencies of the 5 most predominant T cell clonotypes detected within the ITGA1^+^/ITGAE^+^ population (right barplot), in the ITGA1^neg^/ITGAE^neg^ (left barplot) and ITGA1^+^/ITGAE^neg^ (middle barplot) populations for each patient. Each square represents the frequency of a clonotype in the corresponding population. (B) Fold change of the most predominant clonotypes within the ITGA1^+^/ITGAE^+^ population. Dots represent the 5 most predominant T cell clonotypes detected in the ITGA1^+^/ITGAE^+^ population observed using the log2 fold change in ITGA1^+^/ITGAE^+^ (right) and ITGA1^+^/ITGAE^neg^ (middle) compared with the ITGA1^neg^/ITGAE^neg^ (left) cell populations by patient. Each dot is linked across cell populations by a line colored by patient. Statistical analysis was performed using paired Student’s t-tests (****P < 0.0001, **P < 0.01). (C) Jaccard overlap among repetoires was analyzed by generating a heatmap of the Jaccard dissimilarity index calculated across the 3 cell populations: ITGA1^neg^/ITGAE^neg^ (white), ITGA1^+^/ITGAE^neg^ (gray), and ITGA1^+^/ITGAE^+^ (black). The Euclidean distance was used for hierarchical clustering as a color-coded matrix ranging from 0 (minimum dissimilarity) to 1 (maximum dissimilarity). (D) Morisita horn overlap among repetoires was analyzed by generating a heatmap of the Morisita horn dissimilarity index calculated across the 3 cell populations: ITGA1^neg^/ITGAE^neg^(white), ITGA1^+^/ITGAE^neg^ (gray), and ITGA1^+^/ITGAE^+^ (black). The Euclidean distance was used for hierarchical clustering as a color-coded matrix ranging from 0 (minimum dissimilarity) to 1 (maximum dissimilarity). Patients included in this figure are color-coded as patient 1 (orange), patient 2 (green), patient 3 (blue), and patient 4 (red).

### 4. Distribution and infiltration of lung tumors by resident memory CD8^+^ T cell subpopulations

*In situ* multiplex immunofluorescence labeling has enabled us to compare infiltration by different T_RM_ subpopulations either in the tumoural or stromal zone (Fig. 4A). We also included the TCF-1 marker when conducting this staining (Fig. 4A), as it is often considered a marker of stemness potentially associated with response to immunotherapy (Sade-Feldman, Yizhak et al., 2018). We observed a higher density of total CD8^+^T cells, as well as all T_RM_ subpopulations, in the stroma compared to the tumor zone (Fig. 4B). In the tumor, the CD103^+^CD49a^+^ CD8^+^ T_RM_ population was more frequently detected (Mean: 20 ± 62 cells/mm^2^) as compared to the CD103^−^CD49a^+^CD8^+^ T_RM_ population (Mean: 9 ± 11 cells/mm^2^), but both populations were also present in the stroma at higher density (Mean: 119 ± 179 cells/mm^2^ for CD103^+^CD49a^+^ CD8^+^ T cells; Mean: 92 ± 84 cells/mm^2^ for CD103^−^CD49a^+^CD8^+^T cells)(Fig. 4B). The CD49a marker has also been reported to be expressed by endothelial cells (Aman & Margadant, 2023) (Fig. 4A). The TCF-1^+^CD8^+^T cell population also infiltrated the tumor microenvironment in these patients, but these cells were only found in the stroma (Mean: 11 ± 15 cells/mm^2^) and not in the tumor nest. CD103^+^CD49a^+^CD8^+^ T cells were not found to express TCF-1, but 13% of the CD103^−^CD49a^+^CD8^+^ T cells expressed TCF-1 (Fig. 4B, C). TCF-1 expression was mainly observed in the stromally localized populations but not in the epithelial tumor islets (Fig. 4B, C).

**Figure 4:**
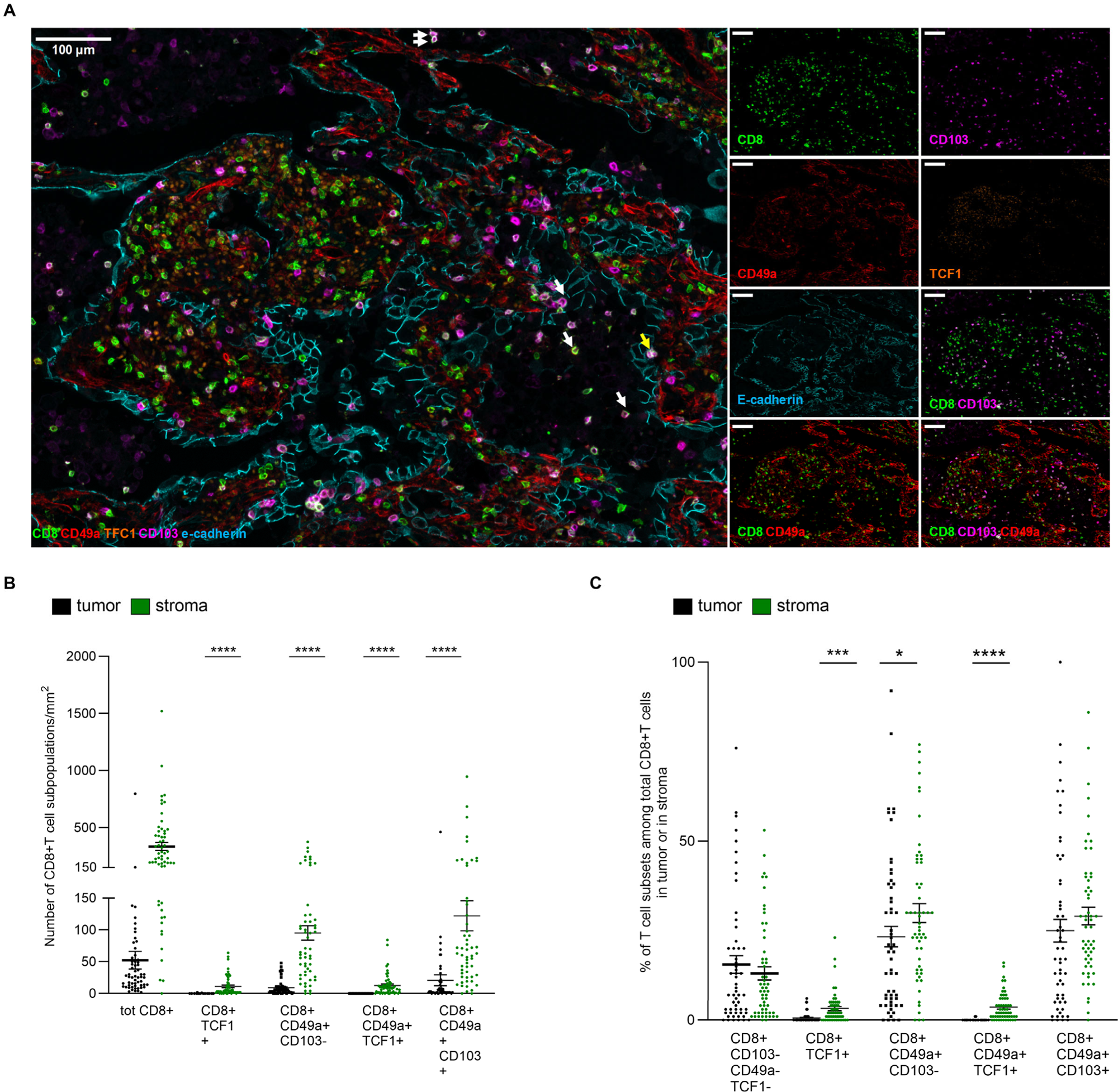
NSCLC tumor infiltration by subpopulations of resident memory CD8^+^ T cells. (A) Representative image of NSCLC tumor infiltration. Multiplexed immunostaining was performed on paraffin-embedded tissues with antibodies to detect CD8, CD103, TCF1, CD49a, and e-cadherin. The InForm^®^ software enabled cell phenotyping and tissue segmentation that was performed by using e-cadherin staining to discriminate between tumor and stromal areas. Automated counting and mapping enabled the phenotyping of T cell subpopulations of non-T_RM_ TILs (defined as CD8^+^CD49a^−^CD103^−^TCF1^+/−^) and of CD8^+^ T_RM_ lymphocytes (defined as CD8^+^CD49a^+^CD103^+^TCF1^−^, white arrow), CD8^+^CD49a^+^CD103^−^TCF1^+^, and CD8^+^CD49a^+^ CD103^−^ TCF1^−^ cells. Original magnification: ×200. Cell numbers (B) and percentages (C) of non-T_RM_ and CD8^+^ T_RM_ were determined via in situ immunofluorescence. Isotype control antibodies were included in each experiment.

### 5. The resident memory CD103^+^CD49a^+^CD8^+^T cell population is the strongest predictor of clinical responses to anti-PD-1 immunotherapy

Cox Proportional-Hazards univariate analysis showed that PD-L1 remained the most predictive biomarker of clinical response (HR = 3.06 [0.002-6.31], P = 0.002) in NSCLC patients undergoing second-line treatment with anti-PD-1(Fig. 5A). Interestingly, the population of T_RM_ CD8^+^ T cells localized in the tumor co-expressing CD103 and CD49a markers was also a pre-treatment feature correlated (HR = 2.41 [1.26-4.62], P = 0.008) with clinical response (Fig. 5A), but this association did not remain true in the stroma (HR = 1.76 [0.092-3.39], P = 0.09). Intratumoural infiltration by CD103^+^CD49a^+^CD8^+^T cells did not predict clinical response as defined by the RECIST criteria (data not shown).

**Figure 5:**
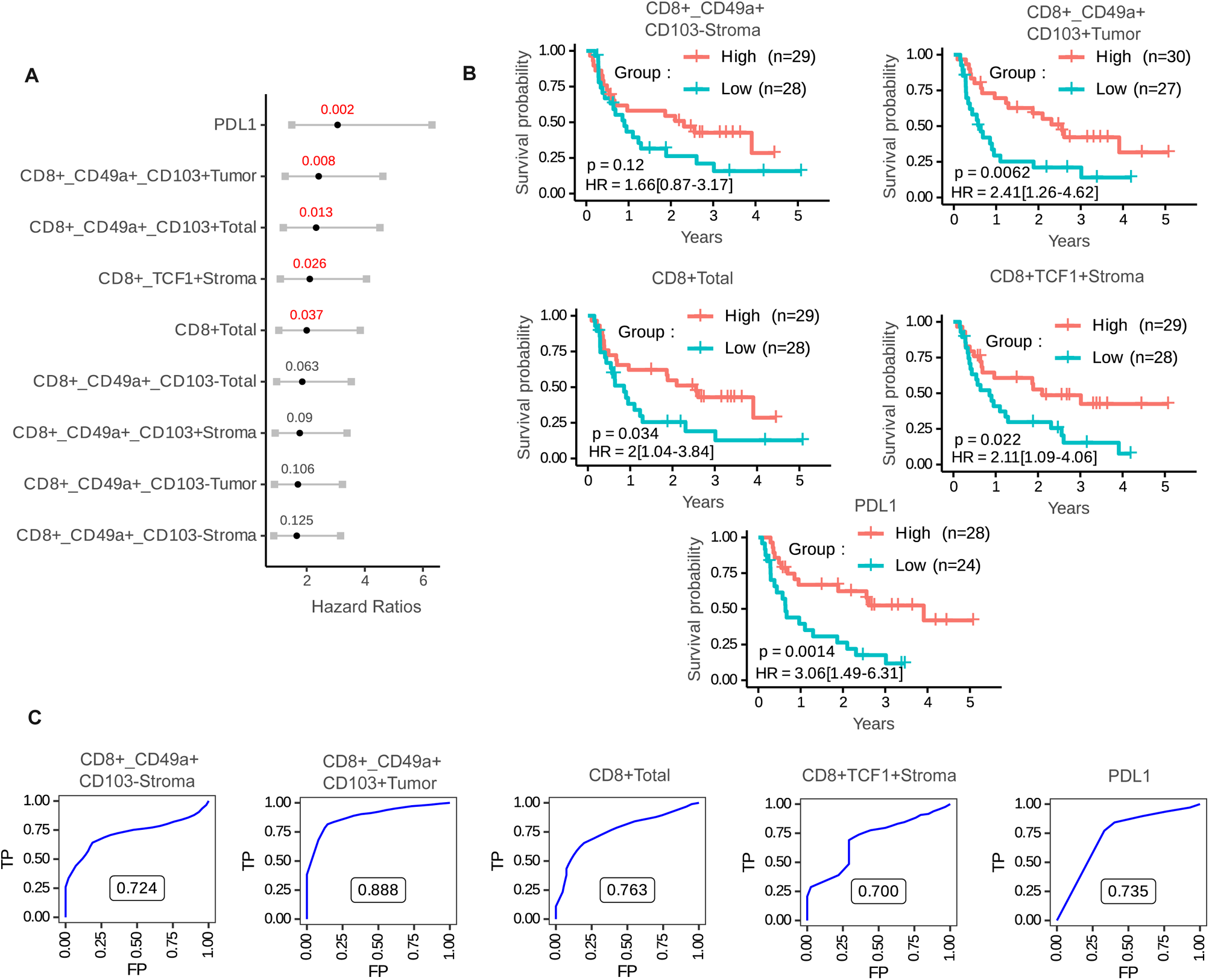
Impact of the infiltration of various subpopulations of CD8^+^T cells in the NSCLC tumor microenvironment on 2^nd^ line therapeutic outcomes. (A) Forest plot showing the Hazard Ratios (HRs) and 95% confidence intervals computed using a univariate Cox model. The infiltration of several subsets of CD8^+^T cells and PD-L1 expression were quantified, using the median as a cut-off for dichotomization. Variables are ordered according to decreasing Wald statistic values. The sublocalization of these subpopulations in the stroma or the tumor or not (total) was taken into account. P < 0.05 was considered significant (in red). (B) Kaplan-Meier curves corresponding to the overall survival of patients with NSCLC grouped according to tumoral or stromal infiltration by subpopulations of resident memory CD8^+^ T cells or total TCF1^+^CD8^+^T cells and the expression of PD-L1 on tumor cells. Each variable was dichotomized separately based on the median value in order to define low and high groups. Log-rank test values are first displayed together with HRs, 95% confidence intervals, and P-values from the Wald test computed using a univariate Cox model. (C) Time-dependent ROC curves were used to analyze the sensitivity and specificity of the two subpopulations of resident memory CD8+ T cells, total CD8^+^T cells, TCF1^+^CD8^+^T cells, and PD-L1 when predicting 2-year overall survival. For each variable, only patients whose variable values were located in the extreme tertiles of the corresponding distribution were included. The resulting area under the curve (AUC) values are shown

In the same analysis, total CD8^+^ T cells (HR = 2.00 [1.04-3.84], P = 0.037) and TCF-1-expressing CD8^+^ T cells (HR = 2.11 [1.09-4.06], P = 0.025) also served as biomarkers associated with clinical response (Fig. 5A). In contrast, the resident memory CD103^−^ CD49a^+^CD8^+^ T cell population, did not predict clinical response to immunotherapy (Fig. 5A) irrespective of its stromal or tumoral location (HR = 1.66 [0.87,3.17], P = 0.12 and HR = 1.70[0.89-3.22), P = 0.106, respectively)

These results were confirmed through Kaplan-Meier survival curve analyses and log-rank tests, confirming the relationships between survival and infiltration by CD103^+^CD49a^+^ CD8^+^T cells, total CD8 and TCF-1^+^CD8^+^ T cells, well as the expression of PD-L1 by tumor cells (TCs) (Fig. 5B). In contrast, only PD-L1 expression by TCs and infiltration by TCF-1^+^CD8^+^ T cells were correlated with PFS (HR = 2.89 [1.52-5.5], P= 0.01 and HR = 1.84 [1.02-3.31], P = 0.043) in these immunotherapy-treated patients (Fig. S7A). This observation was made when comparing two groups of patients dichotomized using the median values for the parameters of interest. When we focused on extreme values using tertiles as cut-offs, the intratumoral CD103^+^CD49a^+^ CD8^+^ cell population appeared to be also prognostic variable for PFS (HR =2.22[1.05-4.67], P = 0.036) (Fig. S7B).

ROC curve analyses of CD103^+^CD49a^+^CD8^+^T cell infiltration yielded an AUC of 0.88 when predicting overall survival at 2 years, with this being the most robust predictor as compared to the other analyzed parameters (Fig. 5C). In a multivariate model, we found that the CD103^+^CD49a^+^CD8^+^ T cell population remained predictive of survival when the model was adjusted for potential cofounders including PD-L1 expression and the infiltration of TCF-1^+^CD8^+^T cells (Table 1).

**Table 1 :** Multivariable model adjusted for PD-L1 level and total stromal TCF-1^+^CD8^+^ T cells. Using Cox proportional Hazard Model, the value of intratumoral CD49a^+^CD103^+^CD8^+^T cells (divided in tertile) in predicting overall survival was evaluated in a multivariate analysis adjusted for PD-L1 and total TCF-1^+^CD8^+^T cells as continuous variables

| Variable | N | HR | 95%CI | P |
| --- | --- | --- | --- | --- |
| TCF-1<sup>+</sup>CD8<sup>+</sup> Stroma | 52 | 0.98 | (0.96,1.00) | 0.12 |
| PD-L1 | 52 | 0.99 | (0.98,1.00) | 0.08 |
| CD49a<sup>+</sup>CD103<sup>+</sup>CD8<sup>+</sup>T cells (Tumor) | 52 |  |  |  |
| Low Tertile | 14 |  |  |  |
| High+Inter Tertile | 38 | 0.39 | (0.16,0.92) | 0.03 |

To confirm these results in a second cohort, we selected NSCLC patients with PD-L1 expression on >50% of TCs who underwent first-line anti-PD-1 treatment (n = 30) or second-line treatment (n = 6). We found that only intratumoral infiltration by CD8^+^ T_RM_ cells co-expressing CD103 and CD49a was correlated with patient survival (HR = 2.77[1.13-6.75], P = 0.025) (Fig. 6A), confirming the results obtained in the discovery cohort. Kaplan-Meier curves and log-rank tests (P = 0.025) were used to analyze patients dichotomized based on median values (Fig. 6B). Tumor infiltration by the CD8^+^ T_RM_ cell population co-expressing CD103 and CD49a was also the only population correlated with PFS when using this same approach to patient dichotomization (HR = 2.62[1.18-5.81], P = 0.018) (Fig. 6C).

**Figure 6:**
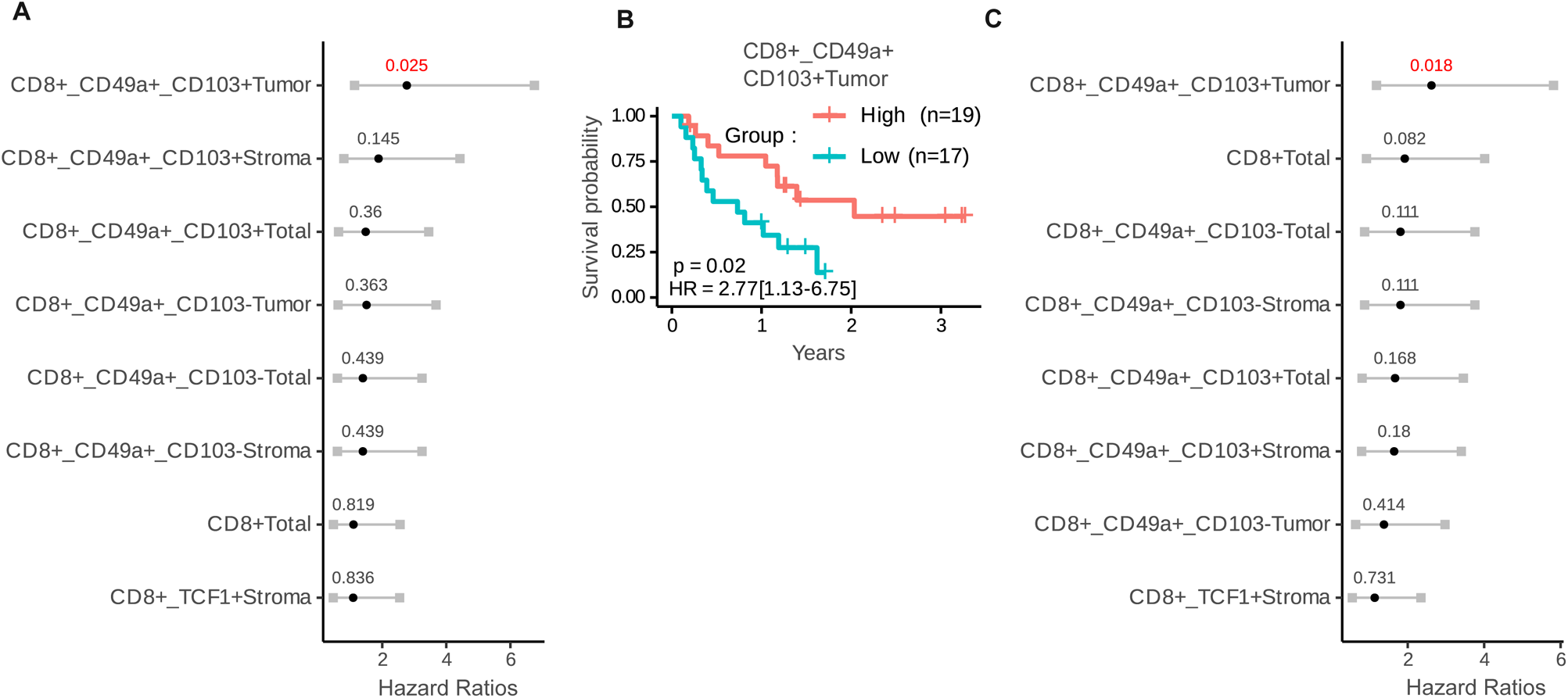
The clinical impact of the infiltration of various subpopulations of CD8+ T cells in the NSCLC tumor microenvironment in a validation cohort. (A) Forest plot representing Cox overall survival regression in NSCLC patients (n = 36). The infiltration of several subsets of CD8^+^ T cells was quantified using the median as a cut-off. The sublocalization of these subpopulations in the stroma or the tumor or not (total) was taken into account. P < 0.05 was considered significant (B) Kaplan-Meier analyses of the overall survival of patients with NSCLC depending on their level of intratumoral CD49a^+^CD103^+^CD8^+^ T cells dichotomized with the median. Statistical analyses were performed with the log-rank test. (C) Forest plot representing Cox progression-free survival regression in NSCLC patients (n = 36). The infiltration of several subsets of CD8^+^ T cells was quantified using the median as a cut-off. The sublocalization of these subpopulations in the stroma or the tumor or not (total) was taken into account. P < 0.05 was considered significant

## DISCUSSION

In this study, we have shown that the intratumoral CD8^+^ T_RM_ cell population co-expressing CD103 and CD49a was highly predictive of response to immunotherapy in two cohorts of lung cancer patients undergoing first- or second-line PD-1 blockade treatment. The predictive performance of this biomarker was also observed in a multivariate analysis. In contrast, the population of CD8^+^T_RM_ cells expressing CD49a without CD103 was not associated with patient response to immunotherapy, irrespective of whether these cells were located in tumors or in the stroma. To the best of our knowledge, no study to date has analyzed the differential prognostic value of resident memory T cell subpopulations. In most reports, CD103 expression alone has been used to define T_RM_ populations. However, different T_RM_ subpopulations have been defined according to a core marker profile that also includes CD49a and CD69. Our CD103^neg^CD49a^+^CD8^+^ T_RM_ population exhibits other T_RM_ characteristics such as CD69 expression, and expresses transcription factors (Runx3 and Hobit) associated with the T_RM_ lineage. Using single-cell analyses, other groups have identified T_RM_ subpopulations in lung cancer at different stages of differentiation that share certain properties with our two populations, but their differential prognostic role has not been reported (Banchereau et al., 2021, Guo, Zhang et al., 2018). Previous work in melanoma had shown that a gene signature corresponding to the CD103^+^CD49^+^CD8^+^T_RM_ cell population was correlated with better overall patient survival (Zitti et al., 2023). The more protective effects of intratumorally rather than stromally localized CD103^+^ expressing T_RM_ observed in this study has also been reported previously in in patients with melanoma, NSCLC, and endometrial cancer (Corgnac et al., 2020, Djenidi et al., 2015, Koh, Kim et al., 2017, Wang, Milne et al., 2016, Workel, Komdeur et al., 2016).

To explain the positive relationship between the CD103^+^CD49a^+^CD8^+^T cell population and overall survival, several hypotheses can be put forward. We have shown in mouse models that CD103^+^CD49a^+^CD8^+^ T cells are more functional than CD103^−^CD49a^+^CD8^+^ T cells, even though they express higher levels of inhibitory receptors such as PD-1 in both humans and mice. Other work has also shown that the CD103^+^ T_RM_ population is superior to its CD103^−^ counterparts in the production of IFN-γ and TNF-α (Lin, Zhang et al., 2020, Watanabe, Gehad et al., 2015). The CD103^+^CD49a^+^CD8^+^T_RM_ population in the epidermis has also been reported as being the most cytotoxic population (Cheuk et al., 2017).

It may seem paradoxical that the most exhausted population of T_RM_ coexpressing CD103 and CD49a is the most functional and associated with a better prognosis. While some studies correlate exhaustion with poor clinical outcomes (Datar, Sanmamed et al., 2019, Ma, Zheng et al., 2019, Mazzaschi, Madeddu et al., 2018, Sade-Feldman et al., 2018), others have defined a subpopulation of CD103^+^CD8^+^T_RM_ cells that appear exhausted, but are also characterized by a proliferative signature and clonal expansion, as well as superior functionality, and are associated with good patient outcomes (Amsen, van Gisbergen et al., 2018, Bassez et al., 2021, Clarke, Panwar et al., 2019, Gueguen, Metoikidou et al., 2021, Hornburg, Desbois et al., 2021, Thommen, Koelzer et al., 2018). To explain these contradictory results, it is worth noting there are different stages of exhaustion that are not equivalent in terms of functionality (Giles, Globig et al., 2023), and certain markers of exhaustion may also, in some situations, correspond to markers of activation (Cheuk et al., 2017). Furthermore, based on some reports, it is likely that the CD103^+^CD49a^+^CD8^+^T cell population is enriched for tumor-specific T cells. Corgnac et al. have shown that this population exhibits enhanced proliferation and cytotoxicity towards autologous tumor cells and frequently displays the oligoclonal expansion of TCR-β clonotypes (Corgnac et al., 2020). We have also shown that this population expresses high levels of PD-1, which is a marker of tumor-reactive TILs in melanoma (Gros, Robbins et al., 2014). Finally, this CD103^+^CD49a^+^CD8^+^T cell population expressed high levels of CD39 (Fig. 2). The CD103^+^CD39^+^ T cell population reportedly has a stronger reactivity against tumors relative to other CD8^+^ T cell subpopulations (Duhen, Duhen et al., 2018, Simoni, Becht et al., 2018). In contrast, the CD103^−^CD49a^+^CD8^+^T cell population may be associated with TILs that are not engaged with antigen (Melssen, Lindsay et al., 2021). Finally, the absence of CD103 in this CD103^−^CD49a^+^CD8^+^T cell population may affect the persistence of this population in the tumor microenvironment and explain its less significant protective role (Mackay, Rahimpour et al., 2013). In addition, antibody blockade of CD103 or CD103 genetic deficiency results in a reduction in tumor-infiltrating T cells and accelerated tumor progression in mice (Malik, Byrne et al., 2017, Murray, Fuertes Marraco et al., 2016).

Our work shows that there is a relationship between the two CD103^+^CD49a^+^CD8^+^ and CD103^−^ CD49a^+^CD8^+^T_RM_ subpopulations, with the former appearing to be more differentiated than the latter. Indeed, these two populations share various TCRs that are amplified in the CD103^+^CD49a^+^CD8^+^T cell population. In addition, the CD103^−^CD49a^+^CD8^+^T cell population exhibits higher levels of TCF-1 expression, a surrogate for progenitor cells, relative to the CD103^+^CD49a^+^CD8^+^T cell population. Other studies have reported common progenitor populations present in the blood that differentiate into distinct T_RM_ subpopulations within tissues (Gueguen et al., 2021, Mackay et al., 2013). Furthermore, it has been shown that during T cell priming, CD49a expression is induced in the lymph node, whereas CD103 expression is acquired after T cell migration in the lung parenchyma and its expression kinetics are more delayed (Haddadi, Thanthrige-Don et al., 2017, Murray et al., 2016). T_RM_ differentiation in the lung parenchyma depends on the interaction of the T_RM_ progenitor population with a specific DC population expressing CXCL16 and membrane IL-15 (Di Pilato, Kfuri-Rubens et al., 2021). Interestingly, the TCF1-expressing progenitor T cell population has also been associated with response to immunotherapy in many cancers (Kurtulus, Madi et al., 2019, Miller, Sen et al., 2019, Philip & Schietinger, 2022, Sade-Feldman et al., 2018, Siddiqui, Schaeuble et al., 2019). We also found that these stromal TCF-1^+^CD8^+^T cells are able to predict responses to immunotherapy, although in a multivariate analysis, only the intratumoral CD103^+^CD49a^+^CD8 T_RM_ population which does not express TCF-1 remained statistically significant as a predictor of this clinical response.

One of the limitations of this study analyzing and comparing the predictive value of subpopulations of T_RM_ as a parameter of response to immunotherapy is that it was performed against total CD8, TCF-1^+^ progenitor T cells, and PD-L1, but not against markers of spatial interactions between PD-1 and PD-L1 or PD-L1 and CD8. These emerging biomarkers have also been reported to predict response to immunotherapy in NSCLC (Ghiringhelli, Bibeau et al., 2023, Sanchez-Magraner, Gumuzio et al., 2023).

This work provides a better understanding of why the CD103^−^CD49a^+^CD8^+^T_RM_ population induced after systemic vaccination does not inhibit tumor growth, in contrast to the CD103^+^CD49a^+^T cell population induced by i.n. immunization (Nizard et al., 2017). We have previously shown that anti-CD49a antibodies reverse the therapeutic effect of the nasally administered vaccines, but were unable to distinguish which subpopulation (CD103^+^ or CD103^neg^) of T_RM_ expressing CD49a was targeted by the vaccine (Sandoval et al., 2013). Cell transfer experiments were inconclusive (Nizard et al., 2017, Sandoval et al., 2013). This strengthens the rationale for mucosal vaccination to induce this protective T_RM_ subpopulation coexpressing CD103 and CD49a.

In clinical practice, patients with lung cancer expressing >50% PD-L1 are treated with anti-PD-1 alone or in combination with chemotherapy. The quantification of the intratumoral T_RM_ subpopulation co-expressing CD103 and CD49a could help guide clinical decision-making when considering these different therapeutic options.

## METHODS

Sex was not considered as a clinical variable in this study

Investigators have been blinded to the patient clinical outcome during the experiment

### 1. Patient sample collection

Two tumor collections from lung cancer patients were provided for this study: i) A colcheckpoint cohort which included tumor tissues from lung cancer patients treated with anti-PD-1 (pembrolizumab or nivolumab) at the Hôpital Europeen Georges Pompidou (HEGP), regardless of the therapeutic line (1^st^ or 2^nd^ line or beyond). Pre-therapeutic biopsies (less than 6 months before the start of immunotherapy) were preferred for most patients, but archival biopsies were also available for some patients. This collection started in June 2016. The clinical database was available for this cohort through a local data warehouse (Bastien Rance); ii) The CERTIM (Immunomodulatory Therapies Multidisciplinary Study group) collection was created in February 2015 by Pr F Goldwasser (Department of Medical Oncology, Hôpital Cochin) and is a collaborative French multidisciplinary network of physicians involved in oncology and research, based at the Cochin Hospital (Paris, France). For this study, it enrolled lung cancer patients treated with anti-PD-1 agents (nivolumab or pembrolizumab). At least one tumor biopsy was available for each patient included in the CERTIM cohort with a varying length of time before the start of immunotherapy, as well as a clinical database for each patient.

From these 2 collections, we selected a discovery cohort of 57 lung cancer patients who underwent 2^nd^ line treatment with anti-PD-1 and a validation cohort of 36 patients, 30 of whom underwent 1^st^ line anti-PD-1 treatment and 6 of whom underwent 2^nd^ line treatment, and for whom tumor tissue from lung localization was available to avoid bias due to localization to other sites.

Flow chart analyses for these cohorts are shown in Fig S1. Characteristics of these two cohorts are presented in Tables S1 and S2.

### 2. Experimental animals

Wild-type female C57BL/6J mice were purchased from Janvier Labs. Experiments were performed using mice 8-10 weeks of age. All mice were housed in an INSERM U970-PARCC animal facility under specific pathogen-free conditions. Experimental protocols were approved by the ethics committee of Université Paris Cité (CEEA 34; approval MESR29315) in accordance with European guidelines (EC2010/63).

### 3. Murine vaccination and sample preparation

STxB-E7 is a DC target-based vaccine chemically linked to the HPV16 E7_43-57_ antigen as described previously (Karaki, Blanc et al., 2021). Anesthetized mice were immunized twice on day 0 and day 14 via intranasal (i.n.) or intramuscular (i.m,) vaccination using STxB-E7, with alpha-galactosylceramide (α-GalCer) as an adjuvant (Funakoshi, Tebu-bio France). On day 21, mice were sacrificed. Intravascular staining was performed to discriminate between tissue-localized and blood-borne cells as described by Anderson et al. (Anderson, Sung et al., 2012). Briefly, 5 µg of anti-CD8α APC-efluo780 (clone 53-6-7, eBioscience/Thermofisher) was injected intravenously (i.v.) 3 min prior to bronchoalveolar lavage fluid (BAL) and tissue collection. BAL was collected from anesthetized mice by flushing the lungs with PBS-EDTA (0.5 mM) via a cannula inserted into the trachea (5 washes x 1 mL).

Lungs were perfused with PBS-EDTA (0.5 mM) and digested in RPMI-1640 medium containing 1 mg/mL collagenase type IV (Life Technologies/ Thermofisher) and 30 µg/mL DNase I (Roche). Lung cells were dissociated using the GentleMACS (Miltenyi Biotec, France) lung programs 1 and 2, with gentle shaking for 30 min at 37°C between both steps. Then, the obtained single-cell suspensions were filtered through a 70-μm strainer, washed with PB containing 2% FBS, suspended in a 40% Percoll solution, layered over a 75% Percoll solution (Sigma-Aldrich), and centrifuged for 20 min at room temperature (RT) at 600 xg. Cells at the interface layer were collected and washed.

After FcR blocking with CD16/32 Ab (clone 93, ebioscience/Life Technologies), cells were first incubated for 30 min at RT with PE-conjugated H-2 D^b^-E7_49–57_ dextramers (Immudex, Bredevej 2A, 2830 Virum, Denmark). Then, cells were washed and stained for surface molecules for 20 min at 4°C in PBS-2%FCS containing anti-mouse CD8β BUV495 (clone YTS156, eBioscience), CD3 PercpCy5.5 (clone 145 2C11, eBioscience/Life Technologies), CD103 Pacific Blue (clone 2E7, Biolegend), CD49a APC or vioFITC (Miltenyi Biotec), and CD69 (clone H1.2F3, Biolegend). For intracellular staining, after surface staining, cells were permeabilized using the FoxP3/Transcription Factor staining buffer set (eBiosciences) according to the manufacturer’s protocols, after which they were stained with an intracellular mAb specific for Tcf1 (clone FAB8224R, Biotechne). All the cells were labeled using the live/dead cell aqua blue viability dye (Life Technologies). Data acquisition was performed with a BD Fortessa X20 instrument (Becton Dickinson), and data from live single cells were analyzed using the FlowJo Software (Tree Star Inc.). Tissue-localized CD8^+^T cells were defined as CD3^+^CD8α^−^CD8β^+^ cells. The adjuvant C-Di-GMP was purchased from InvivoGen (Toulouse, France).

### 4. Single-cell analyses

#### 4.1 Cell sorting

Fresh human tumors were collected in Hank’s Balanced Salt Solution (HBSS). After being mechanically dissociated with a scalpel, they were digested with DNAse I (30 IU/mL, Roche) and Collagenase D (1 mg/mL, Roche) at 37°C for one hour. Primary cells were then isolated by successive filtration with a 70 μm Falcon cell strainer (BD Falcon) and a 20 μm Falcon cell strainer (BD Falcon) and counted using a Malassez chamber. Obtained cells were then stained with the fixable viability stain FVS 520 (eBioscience), APC-conjugated anti-CD8α (Biolegend), and BV421-conjugated anti-NKP46 (Biolegend). NKP46 was used to eliminate natural killer cells from the analysis during sorting.

#### 4.2 Single-cell capture and RNA sequencing

After sorting, CD8^+^ T cells were processed with the Chromium Single-cell 5′ v2 Library Kit (10X Genomics) based on the manufacturer’s instructions. A total of 5,000 cells were loaded per channel of the 10X Chromium Controller as previously described (Benhamouda, Sam et al., 2022). The instrument initially produces an emulsion of 100,000 droplets containing zero or one cell, a single bead coated with a single 16-nucleotide barcode common to all cyclic DNA (cDNA) generated from the same cell, a UMI (Unique Molecular Identifier) sequence of 10 nucleotides specific to each transcript, a sequencing primer (R1), a poly (dT) sequence and all the reagents necessary for cell lysis and RNA reverse transcription. The emulsion was then gently transferred to a plate and incubated at 53°C for 45 min for reverse transcription, followed by the lysis of the cells.

cDNA was then purified and amplified for 12 cycles and used for library preparation. Single-cell barcoded cDNA libraries were again amplified by PCR after the addition of adaptors. The final libraries were qualified and quantified using Bioanalyzer (Agilent) and Qubit (Invitrogen) instruments, pooled together, and sequenced on the Illumina HiSeq X platform. Cells were sequenced to obtain about 30,000 reads per cell.

#### 4.3 Single-cell RNA sequencing data analysis

The raw deep sequencing data were processed using the 10X Genomics CellRanger software package (v 2.1. or 3.0.2) with the GRCh38-3.0.0 reference genome. Subsequent counts were analyzed using scShinyHub (https://github.com/baj12/scShinyHub) with the number of reads per cell being normalized by dividing by the total number of reads for a given cell and then log2(x+1) transformed and scaled with a factor of 1000. Cell selection was performed manually using scShinyHub.

### 5. T cell receptor (TCR) analyses

#### 5.1 Data Source

The raw data used for this TCR repertoire analysis came from the 4 fresh lung tumors described above that were used for single-cell transcriptomic analyses.

#### 5.2 Data pre-processing

Single-cell immune profiling raw data were processed using Cell Ranger (v 6.1.1) using – reference = vdj_GRCh38_alts_ensembl-5.0.0.

#### 5.3 Overlap analysis

Jaccard and Morisita-Horn distances were used to assess the dissimilarity between repertoires (Valkiers, de Vrij et al., 2022). The Jaccard index is a measure of the intersection between two populations relative to the size of their union and is independent of relative abundances, whereas the Morisita-Horn index considers the relative abundance of species in the sample. Both indices vary between 0 (No dissimilarity) and 1 (complete dissimilarity). JSI and Morisita Horn distances were calculated using the vegan package (Oksanen J, Simpson G, Blanchet F, Kindt R, Legendre P, Minchin P, O’Hara R, Solymos P, Stevens M, Szoecs E, Wagner H, Barbour M, Bedward M, Bolker B, Borcard D, Carvalho G, Chirico M, De Caceres M, Durand S, Evangelista H, FitzJohn R, Friendly M, Furneaux B, Hannigan G, Hill M, Lahti L, McGlinn D, Ouellette M, Ribeiro Cunha E, Smith T, Stier A, Ter Braak C, Weedon J (2022). _vegan: Community Ecology Package_. R package version 2.6-4, <https://CRAN.R-project.org/package=vegan)

#### 5.4 Statistical comparisons

R v4.2.3 (www.r-project.org) was used to conduct all statistical analyses. Figures were generated using the ggplot2 package (https://doi.org/10.1007/978-3-319-24277-4), with the exception of heatmaps, which were generated with the pheatmap package (Kolde, R. pheatmap: pretty heatmaps. https://cran.r-project.org/web/packages/pheatmap/index.html (2019). Both indices are presented as dissimilarity values and vary between 0 (no dissimilarity) and 1 (complete dissimilarity).

### 6. Data Availability

Single-cell RNA sequencing (scRNA-seq) data from 4 fresh lung cancer samples were uploaded to the NCBI Gene Expression Omnibus (GEO) archive platform (https://www.ncbi.nlm.nih.gov/geo/) under accession number GSE160243.

### 7. Flow cytometry analyses of TILs

Freshly resected lung tumors and adjacent healthy lung tissue samples obtained from the Institut Mutualiste Montsouris and the Hôpital Marie-Lannelongue were immediately cut into small fragments and digested for 40 min at 37°C using a tumor dissociation kit (Miltenyi Biotech). The dissociated samples were smashed on 100 µm cell strainers, washed, and red blood cell lysis was performed. The recovered single-cell suspension was used for phenotypic analyses performed by direct immunofluorescence with a panel of fluorochrome-conjugated antibodies. Anti-CD3-Alexa700 (UCHT1), anti-CD8-PacificBlue (RPA-T8), anti-CD69-APC-Cy7 (FN50), anti-granzyme-B-FITC (GB11) were supplied by BioLegend. Anti-CD103-BV711 (Ber-ACT8) and anti-Hobit-Alexa647 (Sanquin-Hobit/1) were purchased from BD Biosciences. Anti-CD49-PerCPefluor710 (TS2/7) and anti-PD-1-PeCy7 (eBioJ105) were supplied by Thermo Fisher Scientific. Anti-RUNX3-PE (R3-5G4) and CCR7-PeCy7 (3D12) were purchased from BD Pharmingen. Anti-CD45RA-APC and anti-CD39-APC, were purchased from Miltenyi. Cells were fixed, permeabilized (FoxP3 Buffer Kit, eBioscience) and then stained with fluorochrome-conjugated mAbs. Dead cells were excluded using a LIVE/DEAD Fixable UV dead cell stain kit (Thermo Fisher Scientific). Stained cells were analyzed by flow cytometry using a BD FACS Fortessa flow cytometer (BD Biosciences). Data were processed using FlowJo software (Tree Star Inc..) (Corgnac, Lecluse et al., 2021). This study was approved by the Institutional Review Board of Gustave Roussy (Commission scientifique des Essais thérapeutiques [CSET]) and informed consent was obtained.

### 8. Multiplex immunofluorescence staining

T_RM_ cell infiltration was assessed using formalin-fixed paraffin-embedded (FFPE) slides (4 µM-thick sections) stained with a panel that had been developed manually before being automated with a Leica Bond robot (Leica Biosystems, Wetzlar, Germany). Slides were deparaffinized (Bond Dewax Solution, Leica Biosystems) and then rehydrated via immersion in decreasing concentrations of ethanol in distilled water. Slides were then fixed in 4% paraformaldehyde. Heating mediated-epitope/antigen retrieval was performed with Bond TM Epitope Retrieval 2 (Leica Biosystems). Blocking was performed with plant-based protein blocking buffer (Cell Signaling Technology [CST], MA, USA) for 15 min. Primary mAbs directed against selected antigens were diluted in SignalStain® Antibody Diluent (CST) and incubated for 30 min (except for CD49a, 60 min). Secondary antibodies conjugated to horseradish peroxidase (ImmunoReagents, NC, USA) were then incubated on samples for 15 minutes (except for CD49a, 45 min). Finally, immunofluorescence labeling was performed with the CF® Dye Tyramide from Biotium (CA, USA). The slides were then washed and heated to remove non-adsorbed antibodies/dye, followed by saturation, labeling with the primary and secondary antibodies, repeating this as many times as necessary to achieve multiplexed labeling. The specificity of each antibody was validated using an isotype control.

The list of antibodies and reagents is provided in Table S3. After the final labeling, samples were stained with DAPI for nuclear counterstaining (PerkinElmer, MA, USA) and mounted in EverBrite Mounting Medium (Biotium).

### 9. Multispectral imaging and phenotyping

Biopsies were whole-slide scanned using the Vectra System (PerkinElmer) at 20x magnification. Ten regions of interest were then selected using Phenochart whole-slide reviewer (PerkinElmer). A spectral library enabling the unmixing of dye was prepared using unstained and single-stained tonsil tissues. The inForm Cell Analysis software (PerkinElmer) was used to facilitate cell segmentation and phenotyping based in part on DAPI staining. A phenotyping step was then performed by training the software to recognize cells depending on their expressed surface biomarkers in order to define an analytical algorithm. Cells were then manually checked until the automatized recognition by InForm® was consistent with the visual count. Each phenotype image was checked after software analysis. The InForm® software provides a confidence interval for each phenotyped cell. For the final statistical analysis, cells were taken into account only if the given confidence interval was over 55% for the corresponding phenotype. The data was then analyzed using R software and the phenoptrReports package (Akoya).

### 10. *In vitro* stimulation and multiplex cytokine assay

Single-cell preparations were obtained from the BAL and lungs of mice on day 21 after i.n vaccination as described previously. Total CD8^+^ T cells were isolated by magnetic sorting (EasySep™ Mouse CD8^+^ T Cell Isolation Kit, StemCell Technologies), followed by tetramer E7 and T_RM_ marker staining. Then, E7-specific T_RM_ CD103^+^CD49a^+^, CD103^neg^CD49a^+^, and Teff CD103^neg^CD49a^neg^ populations were sorted by flow cytometry and stimulated (10,000 cells/well) with E7_49-57_ peptide (10 µg/mL) for 18 h. Then, supernatants were harvested and a bead-based multiplexed cytokine immunoassay was performed to detect IL-2, IFNγ, Granzyme B, MIP1a/CCL3, MIP1b/CCL4, and RANTES/CCL5 (R&D Biotechne) according to the manufacturer’s protocol and analyzed using the Bio-Plex 200 platform (Bio-rad). Analyte concentrations were calculated using a standard curve (5 PL regression) with the Bio-Plex manager software.

### 11. Statistical analysis

All statistical analyses were performed using R v 3.4.2. Kaplan–Meier curves (to visualize survival probabilities) and Log-rank tests (to test for statistical significance between groups) were performed using the ggsurvplot function of the survimer package. Univariate analyses for both overall survival (OS) and progression-free survival (PFS) were conducted using a Cox Proportional Hazards model implemented in the coxph function of the survival package, retrieving hazard ratios (HRs), 95% confidence intervals, and Wald statistics to address the statistical significance of the model. For each variable of interest, the data were dichotomized into “Low” and ‘High” groups according to the median or extreme tertile cut-offs of the distribution for the univariate survival analysis.

Time-dependent receiver operating characteristic (ROC) curve and AUC (area under the curve) analyses were conducted with the survivalROC package configured using the Nearest Neighbor Estimation (NNE) method. The Cox proportional hazard model was also used to perform a multivariate analysis to assess whether the prognostic effect of a variable of interest remains significant after adjustment for other potential cofounders. These variables were considered as continuous variables whereas the variable of interest was dichotomized into two groups.

### 12. Study Approval

All clinical investigation have been conducted according to Declaration of Helsinki principles Written informed consent was received from participants prior to inclusion in the study.

The colcheckpoint cohort has received the approval of the CPP Ile de France II (CPP number : 2015-08-04-MS2) and the CNIL declaration (iDP1563364). For the Certim cohort protocol was approved by the local ethics committee (CPP Ile de France II, n°2008-133 and 2012 06-12, MS1) in accordance with article L.1121-1 of the French law.

## Acknowledgments

This work was funded by the Fondation ARC pour la Recherche sur le Cancer (Grant number SIGN’IT20181007747 and PGA 2020 12019110000946_1581 to E. Tartour), the INCA (Institut National du Cancer) (Grant number 2022 PLBIO22-147 to E. Tartour), PCSI 2021 (M2DIA), the Agence Nationale de la Recherche (Labex Immuno-Oncology to E. Tartour), the Institut National du Cancer (Grant SIRIC CARPEM to E. Tartour, L Paolini, S. Oudard), FONCER (to E. Tartour), Fondation pour la Recherche Médicale (FRM) (EQU202103012926 to Ludger Johannes), Institut National du Cancer (INCa) (contract n°2019-1-PLBIO-05-1 to Eric Tartour and Ludger Johannes), La Ligue MucoRNAvax (Convention N°AAPARN 2021.LCC/ChP to Eric Tartour and Ludger Johannes), PEPR RNAvac (ANR-22-PEBI-0007 to Eric Tartour and Ludger Johannes)

We thank the staff of the tumour banks of HEGP (B. Vedie and D. Geromin) for providing the sample materials and the Histology platform of PARCC (C. Lesaffre).

## The Paper Explained

### PROBLEM

Tumor infiltration by resident memory T-CD8 lymphocytes (T_RM_) is often associated with response to immunotherapy, but its predictive value has not yet led to its recommendation in patient management guidelines. There are different sub-populations of T_RM_ whose specific role is not well understood.

### RESULTS

We identified two main T_RM_ subpopulations in tumor-infiltrating lymphocytes derived from non-small cell lung cancer (NSCLC) patients: one co-expressing CD103 and CD49a (DP), and the other expressing only CD49a (MP); both exhibiting additional T_RM_ surface markers like CD69. DP T_RM_ exhibited greater functionality compared to MP T_RM_. Analysis of T-cell receptor (TCR) repertoire and of the stemness marker TCF-1 revealed shared TCRs between populations, with the MP subset appearing more progenitor-like phenotype. In two NSCLC patient cohorts, only DP T_RM_ predicted PD-1 blockade response. Multivariate analysis, including various biomarkers (CD8, TCF1^+^CD8^+^T cells, and PD-L1) associated with responses to anti-PD(L)1, showed that only intra-tumoral infiltration by DP T_RM_ remained significant

### IMPACT

This work shows that it is crucial to distinguish between T_RM_ subpopulations because they have different functionalities, and only the intratumoral T_RM_ population expressing CD103 and CD49a can effectively predict the clinical response of non-small cell lung cancers to immunotherapy.

## Author contributions

LP and TT contributed to the conception of the work, acquisition, analysis, interpretation of the data, and drafting of the manuscript. ET carried out the coordination of the work.

SC conducted experiments, acquired, analyzed and interpreted the data and drafted the manuscript

DD contributed to the conception of the work, analyzed and interpreted the data and drafted the manuscript

JPV, MW, JA contributed to the analysis, and interpretation of the data, as well as critically reviewing the manuscript.

JP, AG, VQ, PB, MH, VL, SM conducted experiments and contributed to analysis and interpretation of the data and review it critically.

LJ, JU, LVV, NG, SdP, PB-R, FG, IC, KL, PLP, HdSB, LG, PR,FMC, EF contributed to analysis and interpretation of the data and review it critically.

All authors participated in final approval of the version to be published.

