## Supplementary material for "Differential predictive value of resident memory CD8^+^T cell subpopulations in non-small-cell lung cancer patients treated by immunotherapy": Table S1-S3 and Figure S1-S7

| Characteristics | Number of patients (%) |
|---|---|
| Median age (n = 57) | 67 |
| Sexe (n = 57)<br>Female<br>Male | <br>21 (36.8)<br>36 (63.1) |
| Histology<br>ADK<br>SCC<br>NA | <br>38 (66.6)<br>14 (24.6)<br>5 (8.8) |
| PD-L1<br>Median<br>PD-L1 > 50 | <br>2<br>17 |
| Tobacco<br>No Smoker<br>Smoker<br>NA | <br>4 (7)<br>48 (84.2)<br>5 (8.8) |
| Survival : Median PFS in days<br>Median OS in days | 99<br>347 |

Table 1 : Characteristics of patients

All patients were metastatic patients treated in 2nd line with anti-PD-1 (nivolumab or pembrolizumab)

| Characteristics | Number of patients (%) |
|---|---|
| Median age (n = 36) | 65 |
| Sexe (n = 36)<br>Female<br>Male | <br>15 (41.7)<br>21 (58.3) |
| Histology<br>ADK<br>SCC<br>Undifferentiated<br>NA | <br>25 (69.4)<br>5 (13.9)<br>2 (5.5)<br>4 (11.1) |
| PD-L1<br>Median<br>PD-L1 > 50 | <br>70<br>65 |
| Tobacco<br>No Smoker<br>Smoker<br>NA | <br>1 (2.7)<br>34 (94.4)<br>1 (2.7) |
| Survival : Median PFS in days<br>Median OS in days | 130<br>405 |

Table2 : Characteristics of patients

All patients were metastatic patients treated in 1st line with anti-PD-1 (pembrolizumab)

(ADK : adenocarcinoma, SCC : squamous cell carcinoma; NA : Not available)

| mAb | Clone | Concentration of primary mAb (ng/ml) | Secondary Ab | TSA Dye |
|---|---|---|---|---|
| **CD8** (Dako, M7103) | C8/144 | N/A | Anti-Mouse UltraPolymer Ab (ImmunoReagent #GAMHRP-050) | CF®680R (Biotium, #92196) |
| **CD103** (Abcam, #129202) | EPR4166(2) | 390 | Anti-Rabbit UltraPolymer Ab (ImmunoReagent #GARHRP-050 | CF®594 (Biotium, #92174) |
| **TCF1** (CST, #2203S) | C63D9 | 400 | Anti-Rabbit UltraPolymer Ab (ImmuoReagent #GARHRP-050 | CF®555 (Biotium, #96021) |
| **CD49a** (Atlas Ab, #AMAb91460) | cl 7217 | 400 | Anti-Mouse UltraPolymer Ab (ImmunoReagent #GAMHRP-050 | CF®488A (Biotium, #92171) |
| **E-cadherine** (CST, 3185S) | 24E10 | 110 | Anti-Rabbit UltraPolymer Ab (ImmuoReagent #GARHRP-050) | CF®430 (Biotium, #96053) |

**Table S3 : List of antibodies selected for multiplex immunofluorescence staining**
Clone and concentration of primary antibodies , reference of secondary antibodies conjugated to horseradish peroxidase and those of CF® Dye Tyramide from Biotium

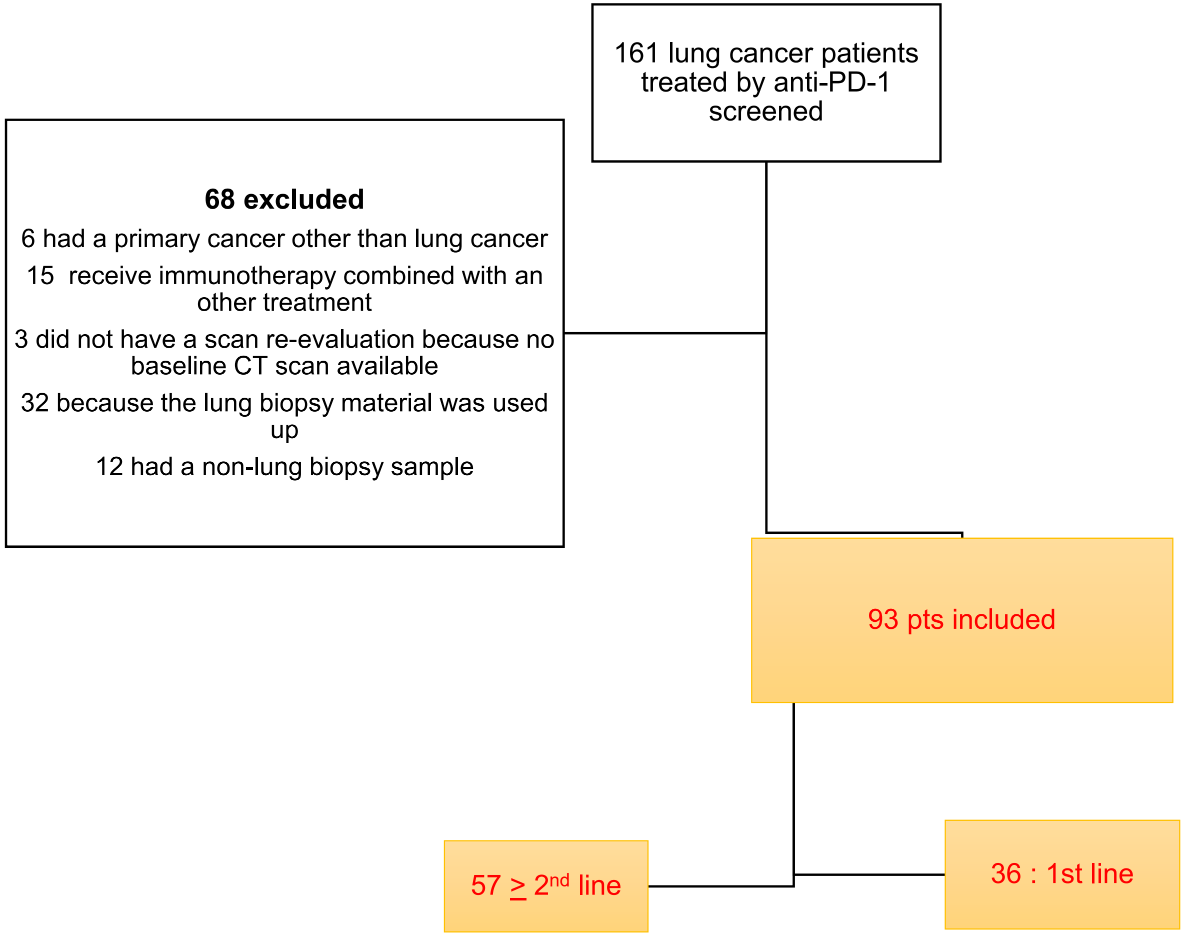


**Fig S1: Flow chart for patient inclusion.**
The numbers of patients assessed for eligibility, causes of exclusion, and patients included in the discovery cohort (n = 57) and the validation cohort (n = 36)

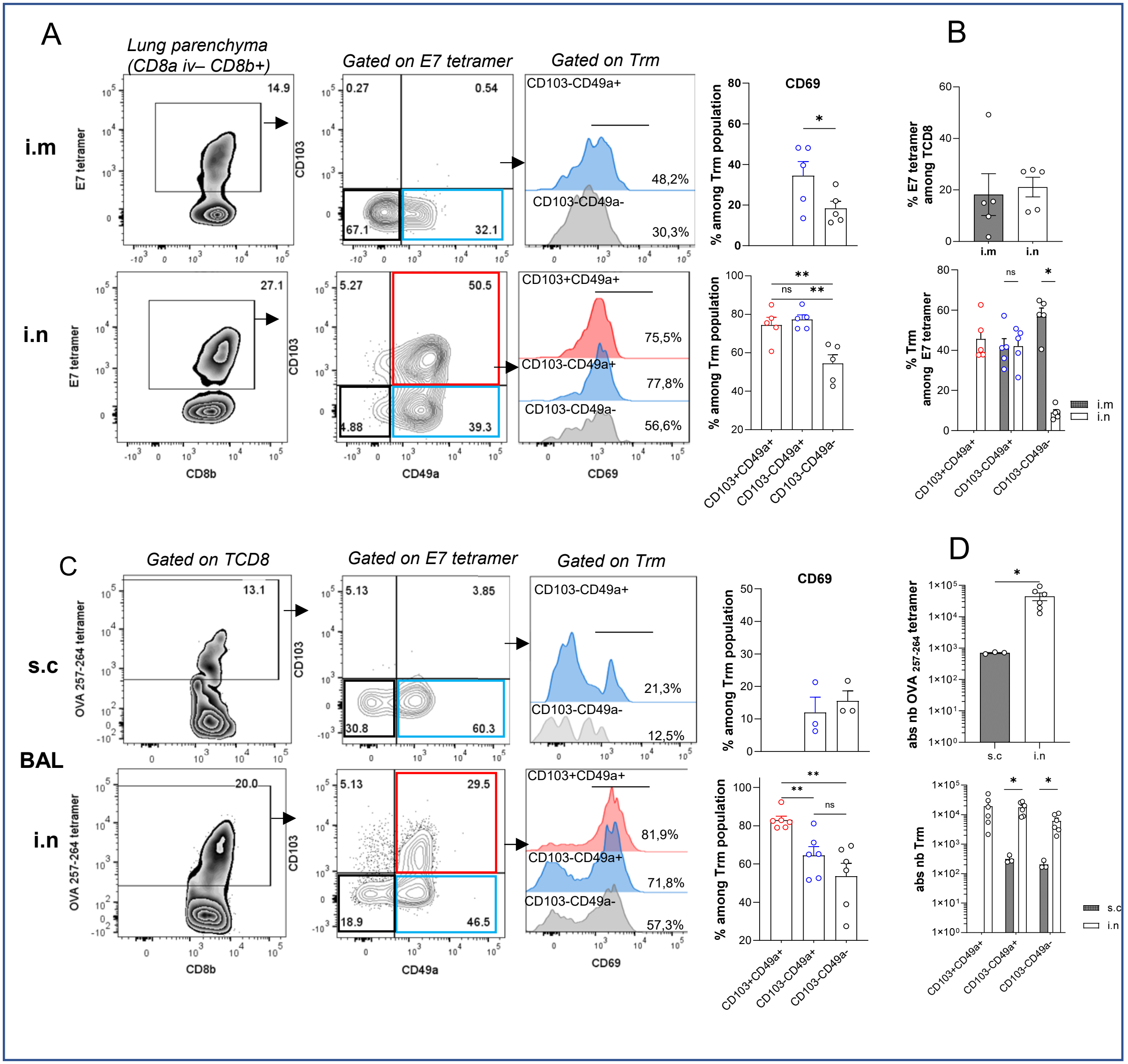


**Fig S2 : Induction of specific Trm CD103$^+$CD49a$^+$CD69$^+$ in the airway and lung after i.n immunization**

C57BL/6J mice were immunized with STxB-E7 (20ug) and αGalcer (2ug) ( A-B) or with Ovalbumine (100ug) +cdg-gmp (10ug) (C-D) by intra-nasal (i.n) or intramuscular (i.m) or sub-cutaneously (s.c) route at day 0 and 14, then sacrified at day 21. CD8a APCefluo780 antibody (3ug) were injected i.v 5 min before sacrifice to discriminate circulating CD8 cells and resident CD8 cells.

A)Representative flow cytometry plots in the lung parenchyma (CD8a iv$^-$CD8b$^+$) of specific E7-tetramer, Trm CD103$^+$CD49a$^+$, Trm CD103$^-$CD49a$^+$, Teff CD103$^-$CD49a$^-$ , and CD69 frequency among these population. B)Percentage of (top)E7-tetramer CD8+ and (bottom) Trm CD103+CD49a+, CD103-CD49a+ and Teff CD103$^-$CD49a$^-$ in lung parenchyma.

C)Representative flow cytometry plots of specific OVA$_{257-264}$ tetramer, Trm CD103$^+$CD49a$^+$, Trm CD103$^-$CD49a$^+$ and Teff CD103$^-$CD49a$^-$ , and CD69 frequency among these population, in the BAL.D) (top) Absolute number of OVA$_{257-264}$ tetramer CD8$^+$ and (bottom) Trm CD103$^+$CD49a$^+$, CD103$^-$CD49a$^+$ and Teff CD103$^-$CD49a$^-$ in the BAL.

Datas are expressed as mean ±sem. 3-6 mice/group, one representative experiment from 2 independent experiments is shown. Analysis of difference within 2 groups were performed with two-side Mann-whitney t-test or one-way ANOVA paired-test with Tukey multiple comparaison.*p<0,05

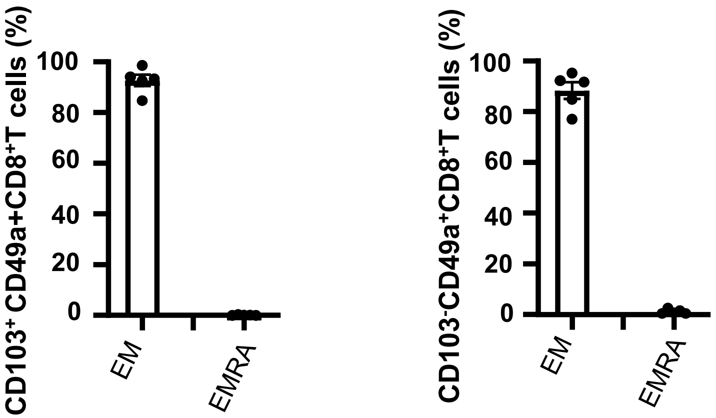


**Fig S3: Percentages of $T_{RM}$ subpopulations among memory effectors and EMRAs.**
Fresh biopsies from patients with NSCLC (n = 3) were dissociated and digested, and flow cytometry analyses of tumor-infiltrating lymphocytes were then performed. The percentages of $T_{RM}$ subpopulations ($CD103^{+}CD49a^{+}$ and $CD103^{-}CD49a^{+}$) among effector (EM) ($CCR7^{-}CD45RA^{-}$) and EMRA ($CCR7^{-}CD45RA^{+}$) cells are shown**.**

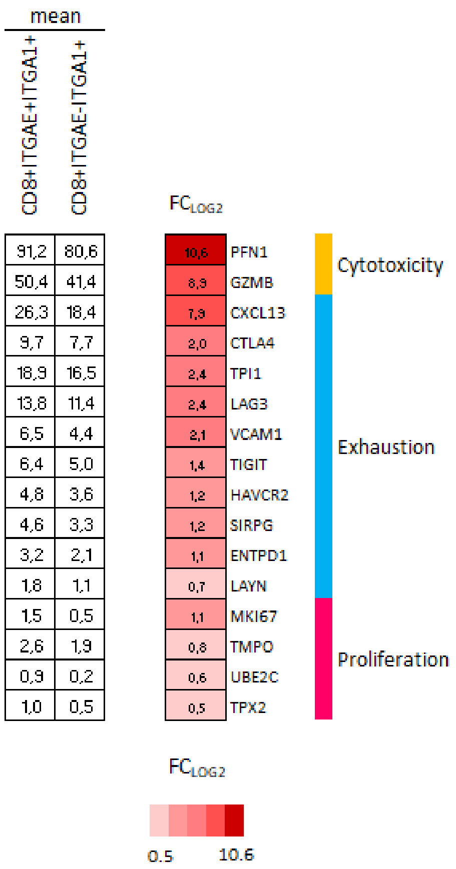


**Fig S4: Genes differentially expressed between $T_{RM}$ populations.**
$CD8^+$ T cells were sorted from 4 fresh lung tumors and submitted for single-cell analyses. The heatmap (right) shows the fold change values for the expression of the differentially expressed genes identified by scRNA-seq between the two $ITGAE^+ITGA1^+$ ($CD103^+CD49a^+$) and $ITGAE^-ITGA1^+$ resident memory $CD8^+$T cells. The mean expression intensities of genes associated with cytotoxicity, exhaustion, and proliferation are shown on the left side of the figure. The difference in the mean expression of these genes between the two $T_{RM}$ subpopulations is expressed as FCLog2 (fold change) in the center of the figure. All of these differentially expressed genes exhibited a P-value < 0.05 after Benjamini–Hochberg correction and a difference in expression of at least 1.4 (FCLog2 = 0.4)

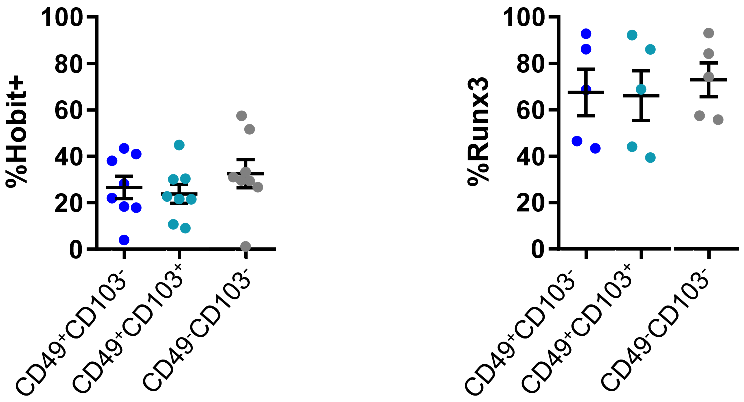


**Figure S5: Expression of Hobit and Runx3 in $T_{RM}$ and Teff subpopulations in TILs derived from lung cancer patients.**

Fresh biopsies from lung cancer patients (n = 5-8) were dissociated and digested. Flow cytometry analyses of TILs were conducted to assess the expression of Hobit and Runx3 for each subpopulation of $CD8^{+}$T cells.

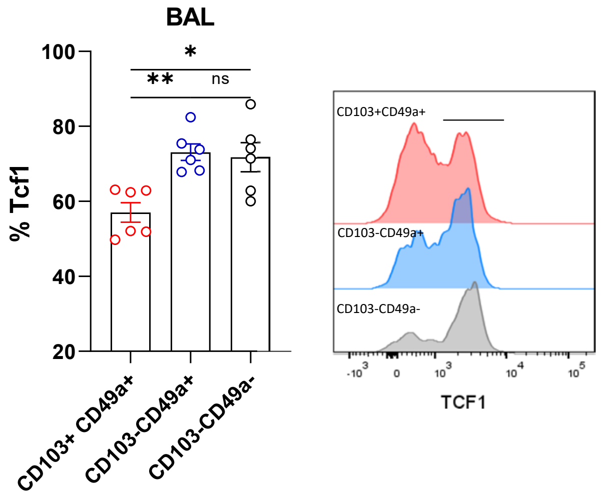


**Fig S6: Expression of the stemness and progenitor marker TCF-1 is decreased on $T_{RM}$ $CD103^+CD49a^+CD8^+$T cells.**

C57BL/6J mice were intranasally immunized with E7 polypeptide and aGalcer on day 0, and were sacrificed on day 9. CD8a APCefluo780 (5 µg) was injected (i.v.) 5 min before sacrifice to discriminate between circulating CD8 cells and resident CD8 cells.

*(left)* Percentage of TCF-1 among E7-specific $T_{RM}$ $CD103^+CD49a^+$, $CD103^-CD49a^+$ and Teff $CD103^-CD49a^-$ in the BAL.

(*right*) Representative flow cytometry histogram of TCF-1 expression among specific E7-tetramer-positive $CD103^+CD49a^+$ $T_{RM}$ , $CD103^-CD49a^+$ $T_{RM}$, and Teff $CD103^-CD49a^-$ in the BAL.

Data are means ± SEM. One representative experiment with 6 mice from 2 independent experiments is presented. Data were analyzed using paired one-way ANOVAs with Tukey's multiple comparison test. *P < 0.05, **P < 0.01.

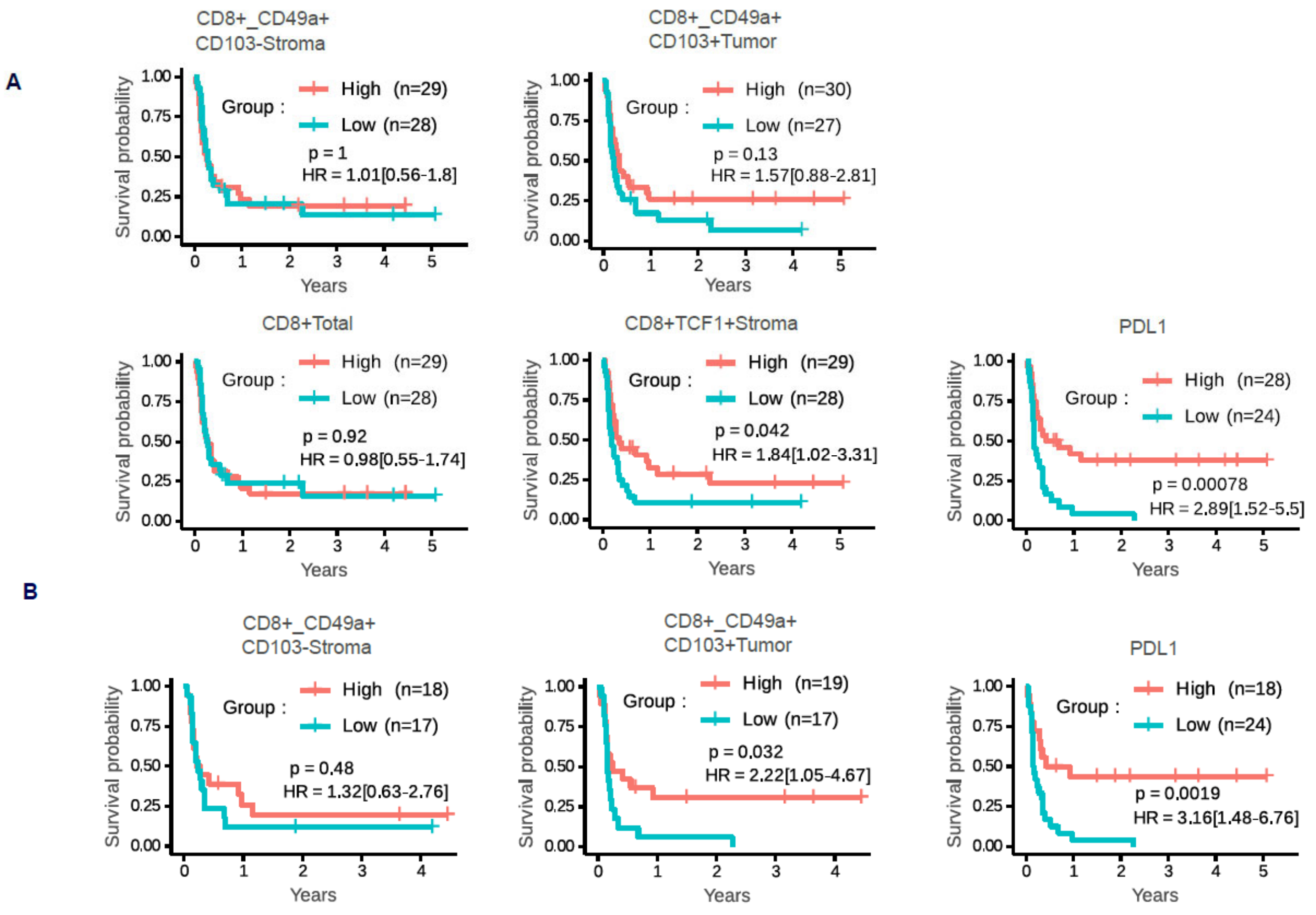


**Fig S7: Analyses of NSCLC patient progression-free survival based on the infiltration of various subpopulations of CD8⁺T cells.**

Kaplan-Meier curves were used to analyze the progression-free survival of NSCLC patients (n = 57) depending on intratumoral or stromal infiltration by subpopulations of resident memory $CD8^+$ T cells or total $TCF1^+CD8^+$T cells, as well as on the expression of PD-L1 on tumor cells. Log-rank test values are presented with hazard ratios (HRs), 95% confidence intervals, and P-values from Wald tests computed using a univariate Cox model. Each variable was dichotomized separately based on the median value (A) or its extreme tertiles (B) in order to define “Low” and “High” patient groups.
